# Microbial resistance and resilience to drought under organic and conventional farming

**DOI:** 10.1101/2024.04.17.589021

**Authors:** Elena Kost, Dominika Kundel, Rafaela Feola Conz, Paul Mäder, Hans-Martin Krause, Johan Six, Jochen Mayer, Martin Hartmann

## Abstract

The impacts of climate change, such as drought, can affect soil microbial communities. These communities are crucial for soil functioning and crop production. Organic and conventional cropping systems promote distinct soil microbiomes and soil organic carbon contents, which might maintain different capacities to mitigate drought effects on cropping systems. A field-scale drought simulation was performed in long-term organically and conventionally managed cropping systems differing in fertilization and pesticide application. The soil microbiome was assessed during and after drought in bulk soil, rhizosphere, and roots of wheat. We found that drought shifted microbial community structures, affecting fungi more strongly than prokaryotes. Microbial communities associated with crops (i.e. rhizosphere and root) were more strongly influenced by drought compared to bulk soil communities. A drought legacy effect was observed in the bulk soil after harvesting and rewetting. The resistance and resilience of the soil microbiome to severe drought did not significantly differ across the organic and conventional cropping systems, although few individual genera (e.g. *Streptomyces*, *Rhizophagus, Actinomadura*, and *Aneurinibacillus*) showed system-specific drought responses. All cropping systems showed relative increases in potential plant growth-promoting genera under drought. This agricultural field study indicated that fungal communities might be less resistant to drought than prokaryotic communities in cropping systems and these effects get more pronounced in closer association with plants. Organic fertilization or the reduction in pesticide application might not have the ability to buffer severe drought stress and additional farming practices might have to be incorporated to improve drought tolerance in cropping systems.

## Introduction

Drought events are projected to increase in frequency, severity, and duration due to climate change in certain regions of the globe [1]. Drought is one major threat to crop yield and health [2] because drought stress leads to osmotic, oxidative, and low-nutrient stress in plants [3]. However, drought stress not only affects plants but also soil microbial communities [4,5], showing possible legacy effects [6]. Soil microbes evolved different mechanisms to avoid or adapt to drought, including osmolyte accumulation [7], production of exopolymeric substances [8], thickening of cell walls [9], and dormancy [10]. Soil respiration, an indicator of microbial activity, and microbial abundance often decrease under water limitation [11,12]. Many studies report an effect of drought on microbial community composition, often indicating that bacteria are more strongly affected by water limitation compared to fungi [4,5]. Fungi have thick cell walls, osmolytes, melanin, and a large hyphal network [13,14], which can improve their drought tolerance. However, bacteria can become dormant during droughts and mostly live in small pores and microaggregates that only dry out slowly, thereby also being able to tolerate drought events [7,10]. Microaggregates were shown to form under reduced precipitation protecting bacterial communities [15]. Slow-growing oligotrophic bacteria that can maintain growth under nutrient-poor conditions are considered to be better adapted to water-limited conditions compared to copiotrophs that thrive under nutrient-rich and well-watered conditions [16].

The soil microbiome is essential for soil functioning and crop production. They are key for climate regulation, nutrient cycling, plant growth promotion and stress tolerance, disease and pathogen control, and degradation of pollutants [17]. Plant roots are associated with bacterial and fungal communities that are located around roots (rhizosphere), on roots (rhizoplane), or inside roots (endosphere) [18]. Some microbes such as plant-growth-promoting rhizobacteria, arbuscular mycorrhizae, and plant endophytes can improve plant drought tolerance and potentially alleviate negative impacts of drought on crops by, for example, increasing plant osmolyte, abscisic acid, or auxin concentrations, decreasing plant ethylene or production of exopolymeric substances [19–21]. Recent studies showed that microbial communities more closely associated with plant roots (i.e. rhizosphere and endosphere) react more strongly to drought compared to bulk soil communities [4,22]. This phenomenon might be directly related to the effects of plant rhizodeposition on the associated microbes [23]. It is known that rhizodeposition can change in quality and quantity during drought, possibly affecting the root-associated microbiome [24].

Many studies assessing the effects of drought on soil microbes have been conducted in grasslands, greenhouses, or in only one type of cropping system. However, it has been reported that different cropping systems, e.g. under organic or conventional practices, can promote distinct soil microbiomes [25], which might differ in their ability to respond to drought. More resilient and resistant microbial communities are suggested to have greater abilities to maintain soil functions under stress such as drought [26]. Resistance and resilience are defined as the ability to tolerate and recover from disturbances, respectively [26].

Organic, biodynamic, and conventional cropping systems differ in fertilization, pesticide application, and crop rotation. Since no synthetic pesticides and mineral fertilizers are applied in organic and biodynamic cropping systems, fertilization is done with green manure, stacked or composted manure, slurry, and by incorporating legumes into the crop rotation. Systems receiving organic amendments generally show higher soil microbial biomass, enzyme activity, microbial diversity, and activity [25,27,28]. Higher soil organic carbon (SOC) contents have been reported in organic cropping systems due to manure application, revealing higher SOC contents in systems receiving composted manure compared to stacked manure [29]. Increased SOC is considered to increase soil aggregation, porosity, and water retention [30]. Thus, higher SOC contents (i.e. improved soil structure and moisture retention) and enhanced microbial diversity and abundance might have the potential to increase microbial resistance and resilience towards drought [31,32]. However, it is not well understood to which extent organic and conventional cropping systems differ in their capacity to increase microbial resistance and resilience under drought [33].

This study compared the effects of severe summer drought on the microbiome in bulk soil, rhizosphere, and roots in long-term organically and conventionally managed cropping systems. For this, we conducted an on-field drought simulation using rainout shelters during winter wheat cultivation in the DOK long-term field trial in Switzerland, which compares different organic and conventional cropping systems since 1978 [25,29,34]. Previous studies have shown that these cropping systems in the DOK trial differ, among others, in SOC content and microbial community structure [25,29]. Sampling took place three times during the drought period and twice after rewetting to assess the microbial resistance and resilience, respectively.

Based on the current literature, we hypothesized that (i) drought effects will be more pronounced on prokaryotic communities compared to fungal communities and (ii) this effect will increase with increasing proximity to the plant (e.g. stronger in root than rhizosphere than bulk soil). We further hypothesized that the (iii) resistance and (iv) resilience of soil microbes towards severe drought stress will depend on the cropping system and increase with higher SOC contents in the following order: a conventionally managed system exclusively receiving mineral fertilization (low SOC), an integrated conventional system receiving a combination of mineral fertilizer and stacked manure (intermediate SOC), and a biodynamic system fertilized with composted manure and slurry (high SOC).

## Methods

### Experimental design

An on-field drought simulation experiment was conducted in the long-term DOK trial, which has been described in more detail by Krause et al. (2022) [29]. Briefly, the field site is located on a haplic luvisol in Therwil, Switzerland (47°30’9.48“N, 7°32’22.02”E). The trial compares five different organic and conventional cropping systems differing in fertilization and pesticide management since 1978. The average annual precipitation at this field site is 840 mm and the mean annual temperature is currently around 11 °C [29].

Rainout shelters, described by Malisch et al. (2016) [35], were established with rain gutters in mid-November 2021 in three cropping systems (Figure 1). The shelters were placed on one side of the plots and the corresponding rainfed controls were established on the other side (Figure 1). Three out of the five cropping systems included in the DOK trial were selected based on the most contrasting biological, physical, and chemical soil properties as found in previous studies [25,34]. The biodynamically managed system (subsequently referred to as BIODYN) is fertilized with composted farmyard manure and slurry, receiving biodynamic preparations, no chemical pesticides, and managed according to the guidelines of Demeter Schweiz (2019) [36]. The other two systems were managed conventionally, one mixed system receiving a combination of stacked farmyard manure and mineral fertilizers (CONFYM) and one exclusively minerally fertilized system (CONMIN). The conventional systems were treated with herbicides, fungicides, insecticides, and synthetic plant growth regulators (chlormequat chloride and trinexapac-ethyl) according to Swiss regulations [37]. The manure-based systems (BIODYN, CONFYM) represent mixed crop-livestock systems and received organic amendments corresponding to a stocking density of 1.4 livestock units per hectare and year. Winter wheat (*Triticum aestivum* var. Wiwa) was sown mid-October 2021. A detailed timeline of all on-field interventions during the experiment is provided in Supplementary Table 1. In brief, shelters were installed in November 2021 and sheltered plots were irrigated during winter 2022 using watering cans with a total of 55 mm of either precipitation or tap water until beginning March. The sheltered plots were then completely deprived from water between 1 April and 14 July 2022. After shelter removal, a rewetting was done on both sheltered and control plots with 36 mm of tap water, and the plots were exposed to rainfed conditions from then on. The entire experiment lasted from mid-November 2021 to mid-September 2022.

**Figure 1:**
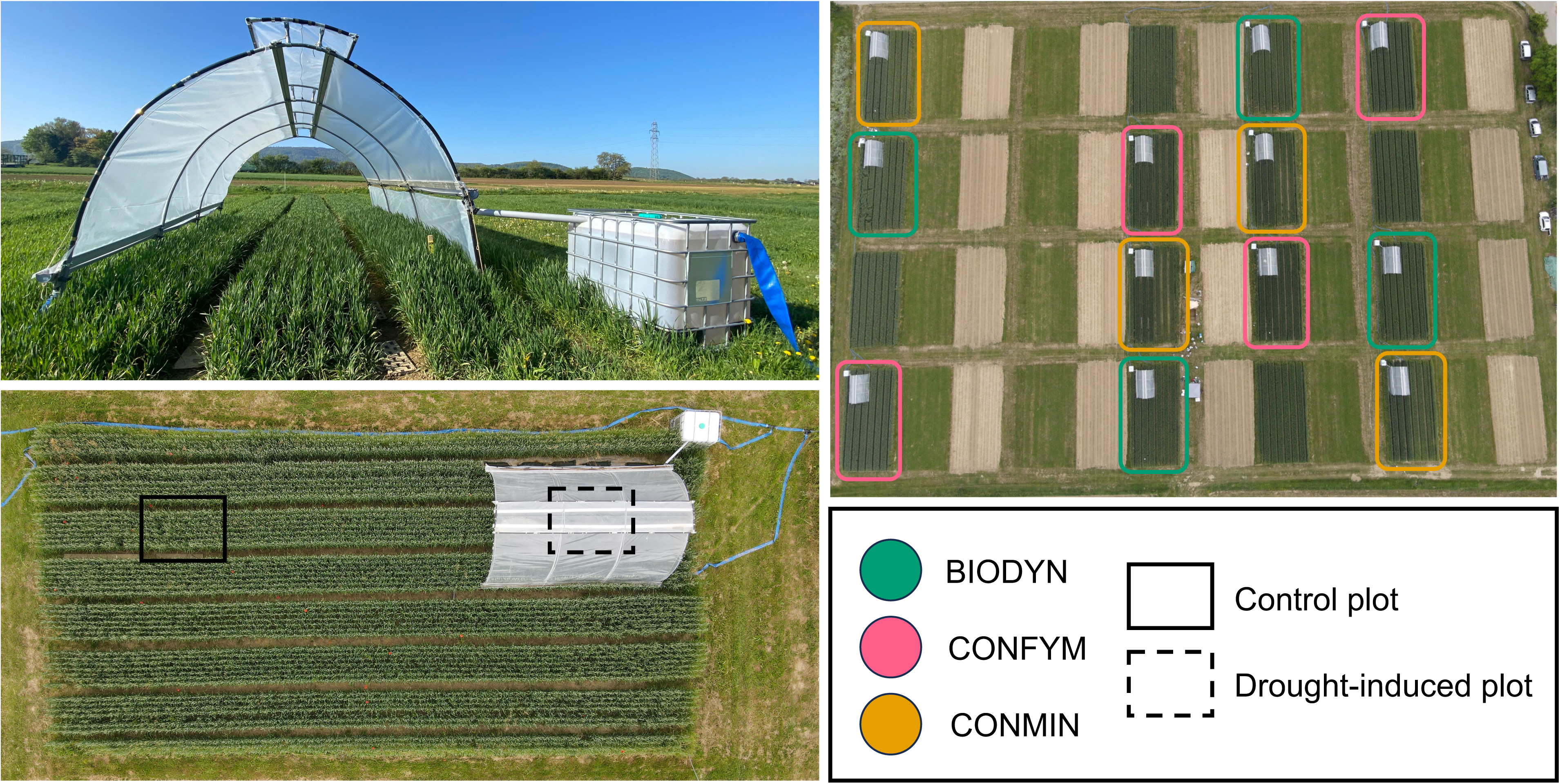
Experimental design of the on-field rainout sheltering experiment in the DOK long-term field trial across three different cropping systems (biodynamic - BIODYN, conventional mixed - CONFYM, and conventional - CONMIN) with winter wheat.

Soil moisture and temperature were monitored in one replication in each of the six experimental treatments at two depths (5 and 20 cm) by time domain reflectometry soil sensors (TDR sensors; METER Group, Pullman, WA, USA) and in all replicated plots by TOMST sensors (TOMST, Prague, Czech Republic) down to 15 cm depth. Gravimetric soil water content (GWC) in 0-15 cm was measured at all sampling campaigns. Air temperature was measured on soil and vegetation level by TOMST and HOBO (EnviroMonitors, Arundel, United Kingdom) sensors, respectively. The latter also measured air humidity. Photosynthetic active radiation (PAR) was measured by PAR Photon Flux Sensors (METER Group) on vegetation level. The HOBO and PAR sensors were installed in the same six plots as the TDR sensors.

### Sampling

Sampling events took place at five timepoints. The first three sampling campaigns were during the wheat growing and drought period at (i) stem elongation, (ii) flowering, and (iii) grain ripening. Plant height, plant, and ear biomass were recorded on an area of 0.042 m^2^ (three wheat rows of 17.5 cm × 8 cm) at each timepoint. Bulk soil samples were taken between the rows with a soil corer (diameter of 5 cm) down to 15 cm (n = 3). Wheat roots with the surrounding soil core were sampled for rhizosphere and root microbiome within rows using a soil auger (diameter of 8 cm) to a depth of 15 cm (n = 3) and loose soil was manually removed by shaking. At the fourth and fifth sampling campaigns (iv) one week and (v) eleven weeks after harvesting and rewetting, respectively, bulk soil was sampled down to 15 cm (n = 3). All bulk soil samples were homogenized and sieved to 5 mm. Bulk soil and root samples were stored at -20 °C until further processing.

### Soil respiration

*In-situ* soil respiration was measured as described in more detail by Barthel et al. (2022) [38]. Briefly, soil respiration was measured weekly during the wheat vegetation period using the non-steady-state, static chamber method with chambers of 30 cm diameter and 30 cm height. Chambers were installed in the field early January. Wheat plants and weeds were removed throughout the seasons within the chambers. For the gas flux measurements, chambers were closed for one hour, and four air samples were collected at 20-minute intervals. Temperature was measured at a metrological station on the field. Carbon dioxide (CO_2_) and methane (CH_4_) concentrations in samples were measured by gas chromatography (456-GC; Scion Instruments, Goes, The Netherlands) using standards covering the expected range of concentrations. The coefficient of determination (R^2^) of the linear regression of 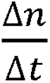 (i.e. the rate of change in concentration in mol s^−1^) from flux data was higher than 0.95 for 94% of the CO_2_ data and 49% of the CH_4_ data.

### Plant and soil measurements

Plant height, plant, and ear fresh weights were recorded in the field. The dry biomass was assessed after drying samples at 40 °C to constant weight. The soil was dried at 105 °C until constant weight to assess the gravimetric water content. The pH was assessed in a soil suspension with deionized water (1:2.5, w/v). Total soil carbon (C) and nitrogen (N) were determined on dried samples with the Dumas method. Magnesium was measured by flame atomic absorption spectroscopy in CaCl_2_ extracts (1:10, w/v). Plant-available soil phosphorus and potassium were measured photometrically and by flame atomic emission in CO_2_-saturated water extract (1:2.5, w/v), respectively.

### Rhizosphere and root separation

After thawing, roots were cut into a 30 mL buffer solution (6.75 g KH_2_PO_4_ and 8.75 g K_2_HPO_4_ in 1000 mL deionized water, adding 200 μL Tween 20 after autoclaving), vortexed for 2 min, and roots were separated into bags. Root samples were freeze-dried and ground with the FastPrep-24™ 5G (MP Biomedical, Irvine, CA, USA). The remaining buffer solution containing the rhizosphere soil was sieved through a 2 mm mesh to remove residual root debris, centrifuged for 10 minutes at 4 °C with 4700 × g, and decanted. The resulting pellet was stored at -20 °C.

### Nucleic acid extraction

The DNeasy ® PowerSoil ® Pro Kit (Qiagen, Hilden, Germany) was used to extract DNA on the QIAcube Connect instrument (Qiagen) according to the manufacturer’s recommendation from 0.25 g homogenized rhizosphere and bulk soil, as well as from 0.04 g homogenized and lyophilized roots. Blanks were included. DNA quality and quantity were assessed via UV/VIS spectrophotometry on a QIAxpert instrument (Qiagen) and normalized to 10 ng μL^-1^.

### Metabarcoding

The bacterial and archaeal (hereafter termed prokaryotic) 16S rRNA gene (V3-V4 region) and the fungal ribosomal internal transcribed spacer (ITS2 region) were PCR amplified with primers 341F/806R and 5.85-Fung/ITS4-Fung using the conditions described in Supplementary Table 2. For root samples mPNA/pPNA clamps (PNA BIO, Newbury Park, CA, USA) were used to inhibit the amplification of organelle DNA with the 16S rRNA gene primers (Supplementary Table 2). PCR products were generated in technical triplicates, which were pooled in equal volumes and sent to the Functional Genomics Center Zurich (FGCZ, Zurich, Switzerland) for indexing PCR. Indexed PCR products were purified, quantified, and pooled in equimolar ratios before pre-sequencing on the Illumina MiniSeq platform (Illumina Inc., San Diego, CA, United States) to inform library re-pooling for optimal equimolarity across samples. Final sequencing was conducted using the v3 chemistry (PE300) on the Illumina MiSeq platform (Illumina Inc.).

The sequence data were quality filtered, delineated into amplicon sequence variants (ASVs), and taxonomically classified against SILVA v138.1 for prokaryotes [39] and UNITE v9.0 for fungi [40] using a customized pipeline largely based on VSEARCH as described previously [41]. The total read number was 14 073 236 (53 920 ± 8969 per sample) for 16S rRNA and 11 725 012 (44 582 ± 16 984 per sample) for ITS sequences. Sequences were assigned to 42 108 and 3801 ASVs after quality control and taxonomic assignment for prokaryotes and fungi, respectively. Prokaryotic ASVs were classified into copiotrophic and oligotrophic lifestyles based on rrn gene copy numbers on the lowest taxonomic rank classified using rrnDB v5.8 [42] and applying the thresholds of ≥ 5 for copiotrophs and < 5 for oligotrophs [43].

### Quantitative real-time PCR

Prokaryotic and fungal abundance in bulk soil and rhizosphere was measured with a SYBR® Green-based quantitative PCR (qPCR) approach targeting the 16S (prokaryotes) or 18S (fungi) rRNA gene as described by Jäger et al. (2023) [44], including a test for potential amplification inhibition, generation of standard curves from purified PCR products of different concentrations, and qPCR amplification of the samples in technical triplicates. The PCR conditions are described in Supplementary Table 2. Amplification efficiencies ranged between 92-100% for (16S) and 75-80% (18S) with an R^2^ of ≥0.95 (16S) and ≥0.99 (18S).

### Statistics

All statistical analyses were performed with R Version v4.3.1 [45] and R Studio Version 2023.06.2+561 [46]. P-values < 0.05 were considered significant unless mentioned otherwise. All permutation-based tests were performed with 9999 permutations. All data was visualized with the R package *tidyverse* version v2.0.0 [47]. Effects of experimental factors on GWC, and plant parameters (height, biomass) were analyzed by a two-way ANOVA when requirements of homogeneity of variance and normal distribution of the residuals were fulfilled. In case the normal distribution of the residuals was not fulfilled, effects of the experimental factors on 16S and 18S rRNA gene copy numbers, the ratio of copiotrophs to oligotrophs, soil chemical properties, soil respirations, and methane were analyzed with a univariate permutational analysis of variance (PERMANOVA) [48] and permutational analysis of multivariate dispersion (PERMDISP) [49] using the *adonis2* and *betadisper* functions in the package *vegan* v2.6.4 [50]. Pairwise comparisons were done with the function *pairwise.perm.manova* in the RVAideMemoire package v0.9-83 [51]. After transforming the logger data (e.g. soil moisture, humidity, PAR, soil and air temperature) using *bestNormalize* v1.9.0 [52], they were analyzed with one-way ANOVA including adjusting for repeated measures.

Rarefaction curves (Supplementary Figure 1) were calculated to inspect the sequencing depth using the *rarecurve* function in *vegan*. To account for differences in sequencing depth across samples [53], ASV tables were 100-fold iteratively subsampled to the minimal read number using the *rrarefy* function in *vegan,* and the average α and β-diversity metrics were calculated based on the 100 subsampled matrices. The Shannon diversity index was calculated using the function *diversity* in *vegan*. β-diversity was assessed based on Bray-Curtis dissimilarities implemented by the function *vegdist* in *vegan*. The effects of experimental factors on α- and β-diversity were assessed by univariate and multivariate PERMANOVA and PERMDISP. Unconstrained ordinations were performed using principal coordinate analysis (PCoA) with the *cmdscale* function in *vegan*. Constrained ordinations were performed using canonical analysis of principal coordinates (CAP) [54] with the *CAPdiscrim* function in the *BiodiversityR* package v2.15.2 [55]. The read counts of each ASV assigned to the same taxonomic group were aggregated across the taxonomic hierarchy and used to test the individual response of taxonomic groups to experimental factors using PERMANOVA followed by adjustments for multiple testing using the *qvalue* function in *qvalue* v2.32.0 [56]. Data were z-transformed for visualization of the differences in relative abundances between all treatments using the *scale* function in R. Genera responding significantly were displayed using iToL v6.8.1 [57], using taxonomic trees built from the taxonomy table using the *taxa2dist* function in *vegan* and the *hclust* function in *ade4* package v1.7-22 [58].

## Results

### Soil and plant measurements

The GWC was significantly reduced during the drought period in sheltered plots compared to the control from on average 26% to 9% (Figure 2), supported by the continuous TOMST and TDR sensor measurements (Supplementary Figure 2). After rewetting, soil moisture increased and showed no significant difference between the water regimes at the second sampling after rewetting (Figure 2). No significant (p > 0.1) interaction was observed between soil water reduction in drought-induced plots and cropping systems at any of the sampling timepoints. Soil temperature below the rainout shelter increased by an average of 15 ± 4% at 5 cm depth and 11 ± 2% at 20 cm depth compared to the control (Supplementary Figure 3). Air temperature slightly increased by 1.2 ± 0.03 °C and 0.4 ± 0.01 °C below the rainout shelter compared to controls assessed at 15 cm above the ground (Supplementary Figure 4) and wheat vegetation level (*F* = 18.4, p = 0.013; data not shown), respectively. Humidity was not influenced by the sheltering (*F* = 0.1, p = 0.782; data not shown), while the mean PAR was reduced by 28 ± 2% due to sheltering (Supplementary Figure 4).

**Figure 2:**
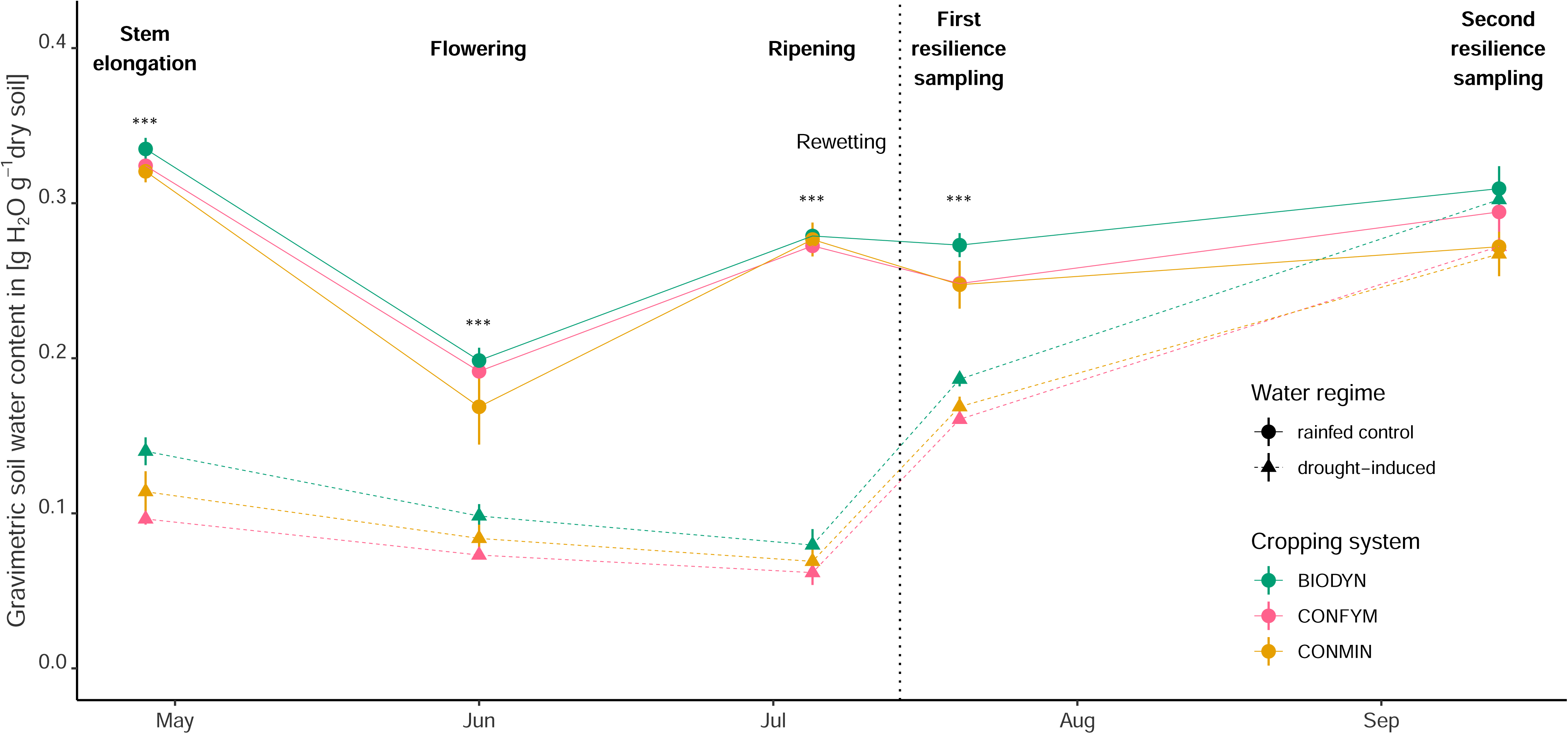
Gravimetric water content (GWC) for each cropping system in drought-induced and rainfed control plots across the five sampling points. Asterisks indicate significant (p < 0.001, n = 12) differences between drought and control plots as tested with ANOVA. Means and standard errors are shown.

Plant-available phosphorus and potassium concentrations were significantly influenced by drought (Supplementary Table 3), showing an increase of 16 ± 5% for phosphorus and 35 ± 14% for potassium during drought in the sheltered plots. The effect of drought on potassium was dependent on the cropping system and increased under drought in all systems but most strongly in the conventional system. This increase in available potassium and phosphorus in the drought plots disappeared after the rewetting (Supplementary Table 4). The other soil chemical properties (i.e. total C and N, plant-available magnesium, pH) showed no significant differences between the water regimes. Cropping systems affected all measured chemical properties during drought and after rewetting (Supplementary Table 3 and 4).

Plant height was significantly increased below the rainout shelters at stem elongation (Supplementary Figure 5). At flowering and ripening, plant height was lower below the shelters compared to the control. However, drought and cropping systems showed an interactive effect, which was reflected by larger differences between sheltered and control plots in the two conventional systems (CONFYM, CONMIN) as compared to the BIODYN system (Supplementary Figure 5). Drought significantly reduced the total fresh weight at flowering and ripening (Supplementary Figure 5). The sheltering significantly increased the total dry biomass of wheat at stem elongation while no differences between the water regimes were observed at flowering and ripening (Supplementary Figure 5).

### Soil respiration

Drought significantly (p < 0.001) reduced *in-situ* soil respiration by an average of 25 ± 8% over the whole drought period, but with strong fluctuations over time (Supplementary Figure 6). Agricultural management significantly influenced soil respiration across both water regimes, having the lowest soil respiration in BIODYN compared to the conventional systems (p < 0.005). The low coefficient of determination of the CH_4_ data indicates that there is no strong methane flux (data not shown). Yet, on average methane uptake was recorded, but with high variability between replicates, nevertheless, showing an increased methane sink by 23 ± 35% under drought compared to the rainfed controls (p < 0.05).

### Microbial abundance

The abundance of prokaryotes and fungi in bulk soil, measured as 16S and 18S rRNA gene copy numbers, were not significantly affected by drought (Supplementary Figure 7). A significant increase of the fungi to prokaryotes (F/P) ratio was found under drought at the first timepoint in the bulk soil and a decrease at the third timepoint in the rhizosphere. There was a significantly lower F/P ratio in BIODYN compared to the conventional systems in the rhizosphere and bulk soil, independent of the water regime (Supplementary Figure 7).

### Microbial diversity and community composition

Since all compartments showed significantly (p < 0.001) distinct microbial communities, compartment data were analyzed separately. Differences in relative abundances of major taxonomic groups between compartments are illustrated in Supplementary Figure 8.

Prokaryotic α-diversity (assessed as Shannon index) was not influenced by drought, whereas fungal α-diversity significantly decreased in the rhizosphere and increased in the root during drought compared to the control (Supplementary Table 5). No interaction between drought response and cropping system on α-diversity was found for fungi or prokaryotes.

PERMANOVA showed that drought significantly affected prokaryotic β-diversity in the rhizosphere and root but not in the bulk soil (Table 1). The effect of drought was stronger in the root (11.5% of the variance explained) than in the rhizosphere (3.1% of the variance explained). Significant differences in fungal β-diversity between drought and control were found for all three compartments, explaining 5.7%, 7.8%, and 6.8% of the variance in bulk soil, rhizosphere, and root, respectively (Table 1). The cropping system had a significant influence on fungal and prokaryotic β-diversity in all compartments, explaining between 10 and 30% of the variance (Table 1, Supplementary Figure 9). The effect of the cropping system decreased from bulk soil (23-31% of the variance) to rhizosphere (18-30%), and root (10-20%). A significant interaction of drought and cropping systems was observed for prokaryotes only in the root, explaining 3.5% of the variance (Table 1). The sampling date explained 1-6% of the variation in β-diversity and significantly affected fungi in all compartments and prokaryotes in the root only (Table 1). The effect of drought depended on the sampling date indicated by a significant interaction in the rhizosphere and root for fungi, and in the root for prokaryotes (Table 1). An increased dissimilarity between the water regimes with proceeding drought was observed mainly for fungi in rhizosphere and root (Supplementary Figure 10).

**Table 1:**
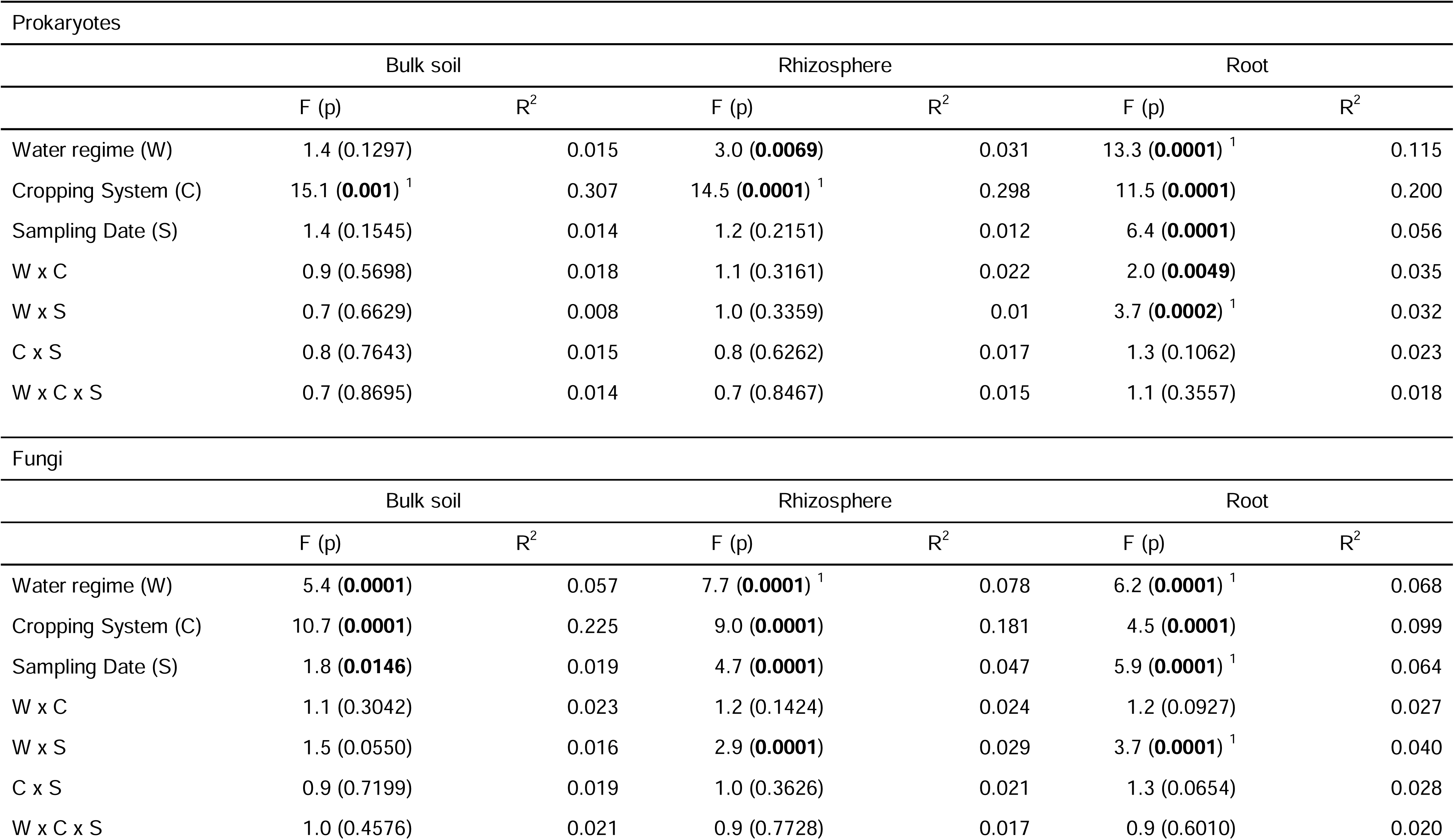
PERMANOVA results (F-ratio, p-value, and R^2^) showing the effect of drought, cropping system, and sampling date on the prokaryotic and fungal β-diversity during the wheat vegetation period. Differences are based on Bray-Curtis dissimilarities and separately analysed for the three compartments (i.e. bulk soil, rhizosphere, and root). Heteroscedasticities are indicated as superscript 1. Values p < 0.05 are indicated in bold.

The CAP using water regime and cropping system as the constraining factors showed distinct clusters between the water regimes during drought in all three cropping systems and in all compartments for fungi and prokaryotes, supported by high reclassification rates (Figure 3). Thus, in contrast to PERMANOVA, CAP and the associated discriminant analysis could resolve differences between water regimes in all compartments and for both communities. In the bulk soil, the cropping system was the main driver of cluster formation (Figure 3 A&B); in the rhizosphere, the two water regimes already showed more distinct clusters (Figure 3 C&D); in the root, the cluster separation was similar between the two water regimes and the cropping systems (Figure 3 E&F).

**Figure 3:**
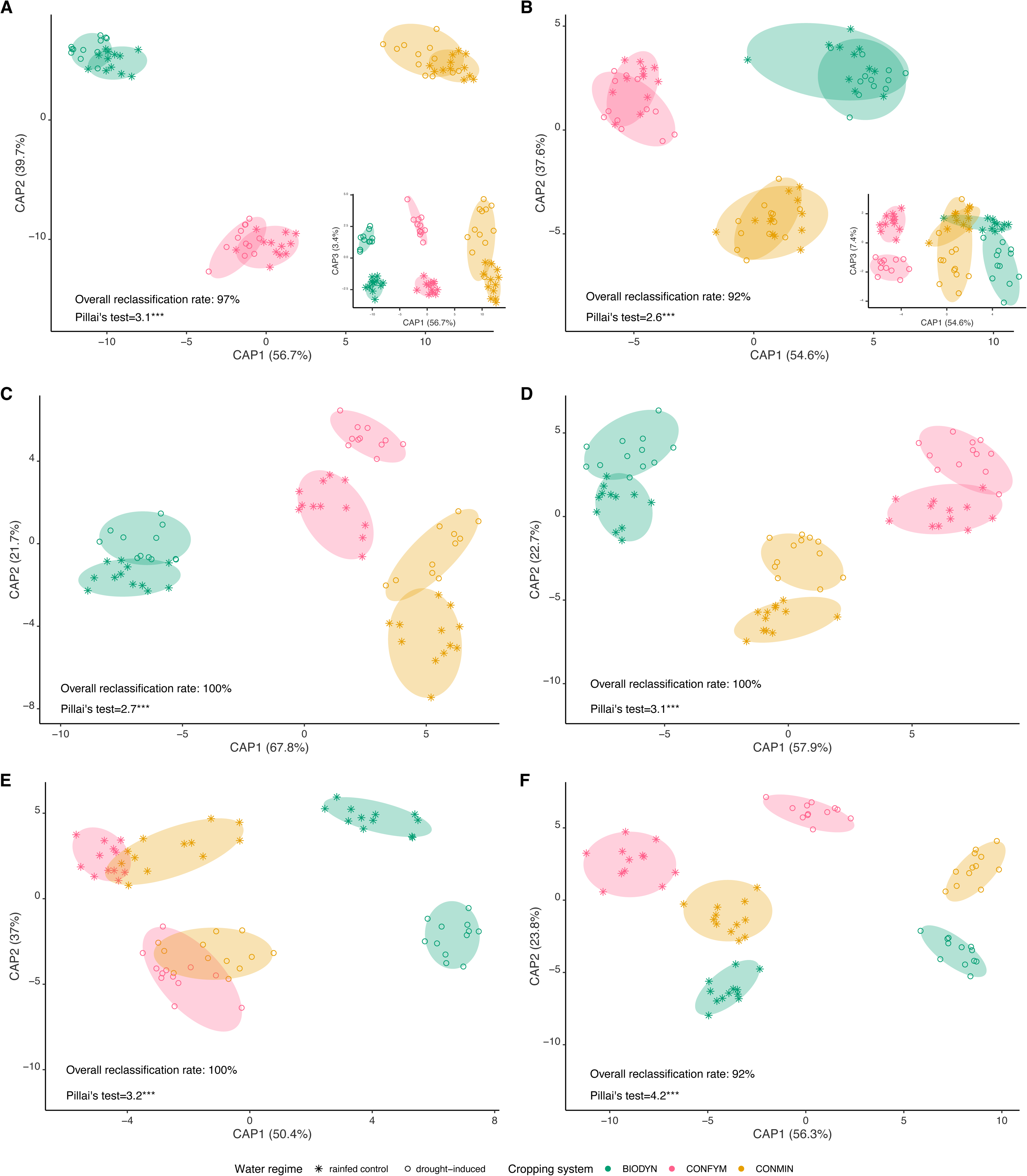
Effects of drought and cropping system on prokaryotic and fungal β-diversity during the drought period. Differences are displayed as canonical analysis of principal coordinates (CAP) maximizing discrimination between water regimes and cropping systems. The CAP overall reclassification rate in percentage, Pillai’s trace statistics, and statistical significance (p < 0.001 ***) are provided in each plot. Panels represent differences in prokaryotic communities in bulk soil (A), rhizosphere (C), and roots (E) as well as fungal communities in bulk soil (B), rhizosphere (D), and roots (F). The amount of between-group variation of each CAP axis is provided in parentheses. For bulk soil, the third dimension is provided to show the separation by the drought treatment.

A CAP for the bulk soil using the water regime as the constraining factor was conducted to evaluate differences in prokaryotic and fungal β-diversity over time (Figure 4). This revealed high reclassification rates for prokaryotes and fungi for both water regimes over the whole experiment (including the period during drought and after rewetting). Prokaryotic and fungal communities at each sampling date separately in each cropping system showed similar reclassification (75-100%) to their respective water regime over the drought period and after the rewetting. In addition, a CAP constraining by water regime and sampling date (whole drought period versus first and second timepoint after rewetting) was performed for each cropping system separately (Supplementary Figure 11). The results showed distinct clusters for fungal and prokaryotic communities at the first and second timepoint after rewetting in the drought-induced treatment compared to the control for all cropping systems, reporting high reclassification rates in all cropping systems. In the control, samples of the drought period and one week after rewetting could hardly be differentiated which was not apparent for the samples from induced drought. PERMANOVA, run for the two sampling dates after rewetting, revealed strong differences in fungal β-diversity and comparatively minor differences in prokaryotic β-diversity between drought-induced and control plots after rewetting. No interactions were reported between the cropping system and water regime after the rewetting (Supplementary Table 6).

**Figure 4:**
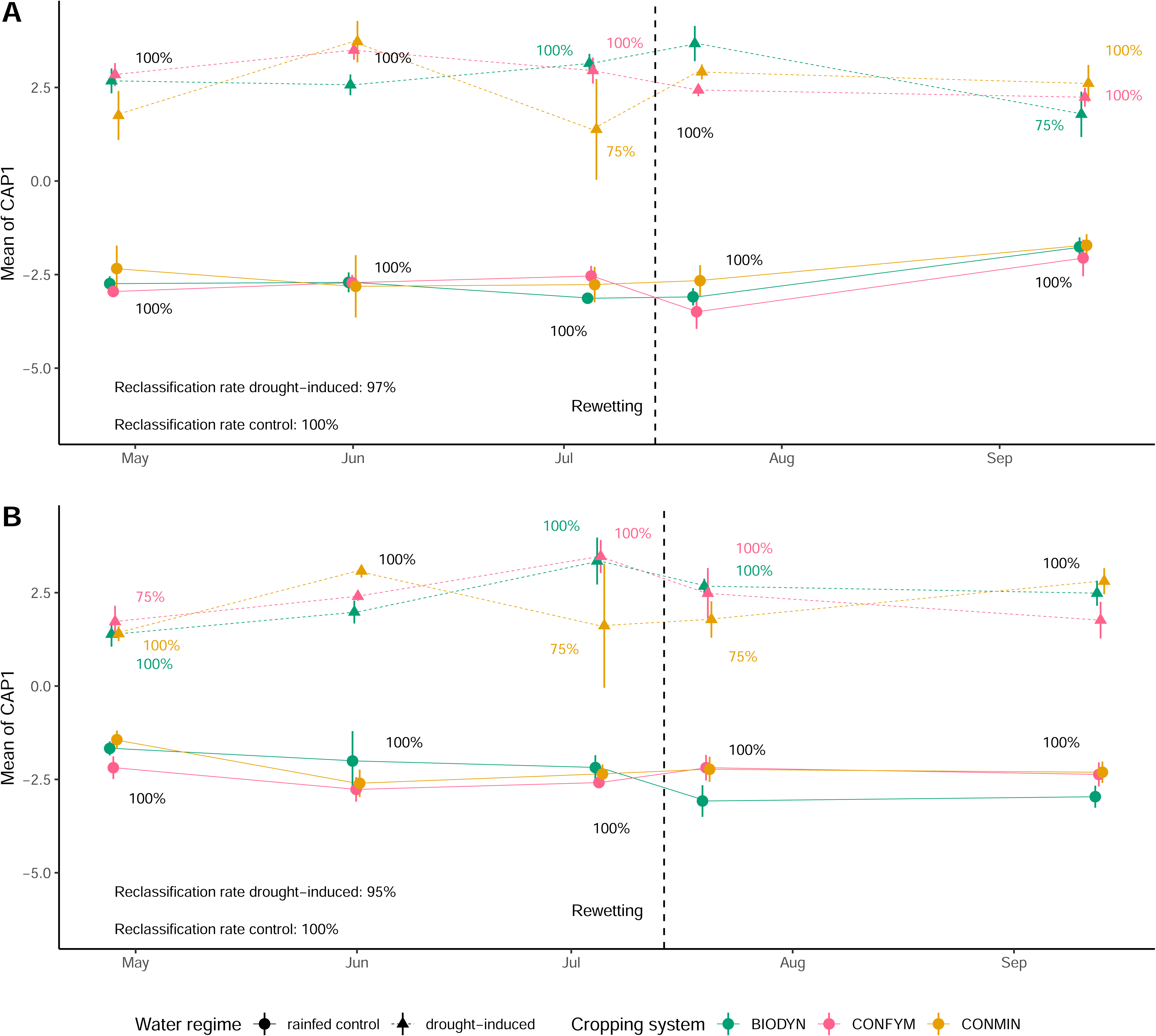
Effects of drought on prokaryotic and fungal β-diversity during drought and after rewetting. Differences are displayed as means and standard errors of the first canonical axis from the canonical analysis of principal coordinates (CAP) maximizing discrimination between water regimes (n = 4). The CAP overall reclassification rate in percentage, Pillai’s trace statistics, and statistical significance (p < 0.001 ***) are provided in each plot. Reclassification rates for each water regime to their water regime at each sampling timepoint and cropping system are provided and displayed in case of differences between cropping systems in the respective color. Panels represent differences in prokaryotic communities (A) and fungal communities (B) in bulk soil. The amount of between-group variation of each CAP axis is provided in parentheses.

### Taxon-level responses to drought

Around 3% (23 out of 696), 13% (91), and 23% (161) prokaryotic genera, and 6% (28 out of 439), 14% (61), and 11% (49) fungal genera were significantly (q < 0.05) altered by drought across all cropping systems in the bulk soil, rhizosphere, and roots, respectively (Figure 5). Genera sensitive to drought were spread over the taxonomic tree, but drought stress tended to increase the relative abundance of genera assigned to *Actinobacteriota* and decrease genera assigned to *Bacteroidota* and *Planctomycetota* in all compartments. In bulk soil, *Cyanobacteria* decreased and *Glomeromycota* increased (Figure 5).

**Figure 5:**
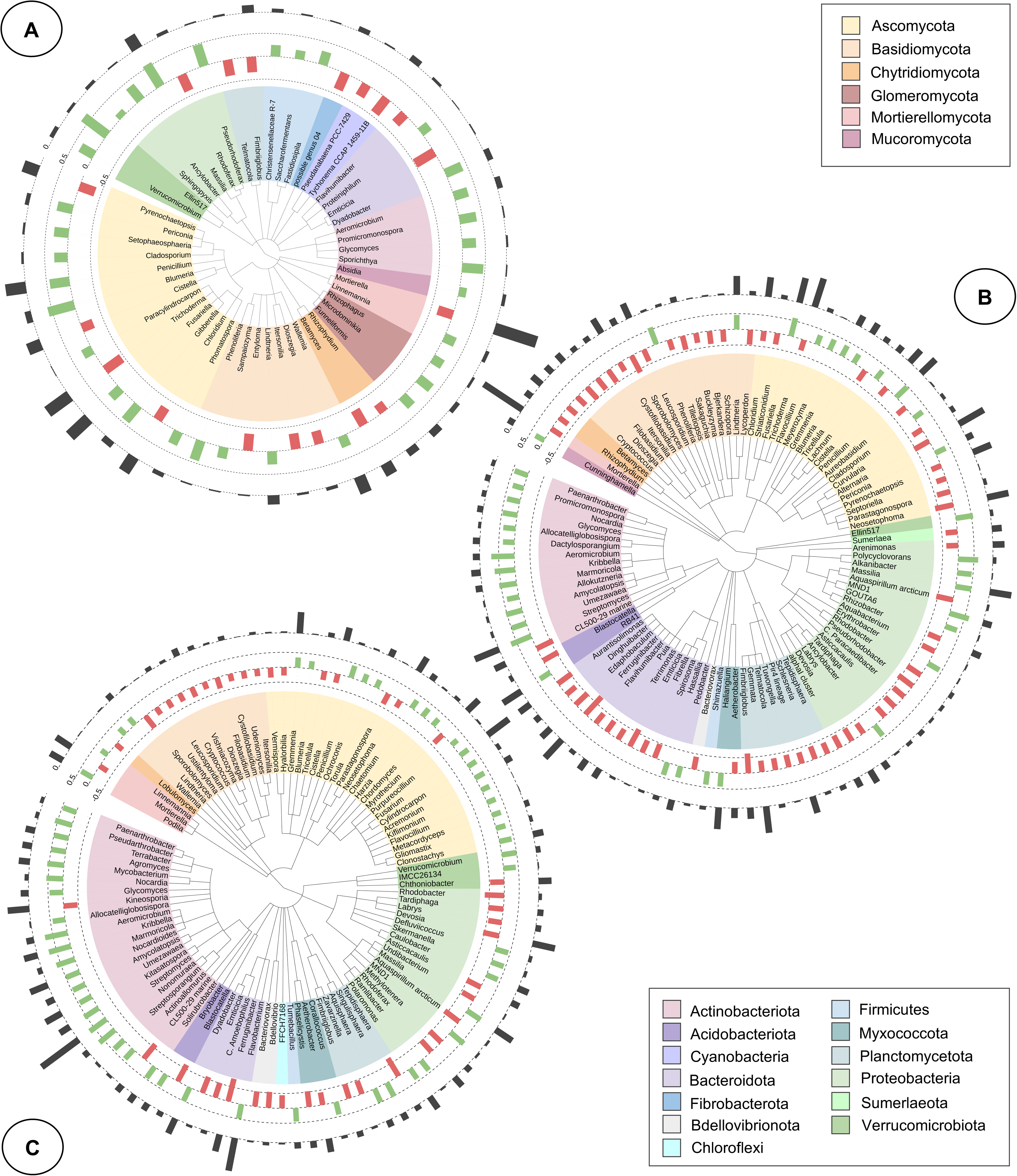
Taxonomic trees displaying prokaryotic and fungal genera in bulk soil (A), rhizosphere (B), and roots (C) responding significantly to drought (PERMANOVA, q < 0.05). All 52 responsive prokaryotic and fungal genera are displayed for the bulk soil, whereas the 60 most strongly reacting prokaryotic and 40 most strongly reacting fungal genera are shown for the rhizosphere and root compartments, respectively. Color ranges indicate corresponding phyla. Colored bar plots showing the z-transformed relative change in abundance of genera either enriched (green) or depleted (red) under drought, respectively. Black bar plots represent the relative square-root transformed mean abundances of genera in the overall community.

Including all compartments, 8% (54 out of 696) of the prokaryotic genera and 5% (20 out of 439) of the fungal genera showed a significant (q < 0.1) cropping system-dependent response to drought (Figure 6). Genera with a cropping system-dependent response to drought in the bulk soil included but were not limited to *Rhizophagus*, *Microdominikia* (both *Glomeromycota*)*, Methanobrevibactera* (*Euryarchaeota*), *Trichococcus, Christensenellaceae R-7, Saccharofermentans, Fastidiosipila, Ercella* (all *Firmicutes*), *Levilinea, Leptolinea* (both *Chloroflexi*), *Roseimarnus, Proteinphilum, Fermentimonas* (all *Bacteroidota*), and *Glycomyces* (*Actinobacteriota*). In the rhizosphere, differentially responsive genera included *Gremmenia, Blumeria* (both *Ascomycota*), *Variovorax, Massilia* (both *Proteobacteria*), *Proteiniphilum* (*Bacteroidota*), *Actinomadura* and *Lechevalieria* (both *Actinobacteriota*). In the roots, differentially responsive genera included for example *Blumeria* (*Asocomycota*), *Paracoccus* (both *Proteobacteria), C. Desulforudis, Sedimentibacter*, *Ruminiclostridium* (all *Firmicutes*), *Solitalea, Proteiniphilum* (both *Bacteriodetes*), *Streptomyces, Kitosatospore, Umezawaea,* and *Salinispora* (all *Actinobacteria*). Results on other taxonomic levels can be found in Supplementary Data 1.

**Figure 6:**
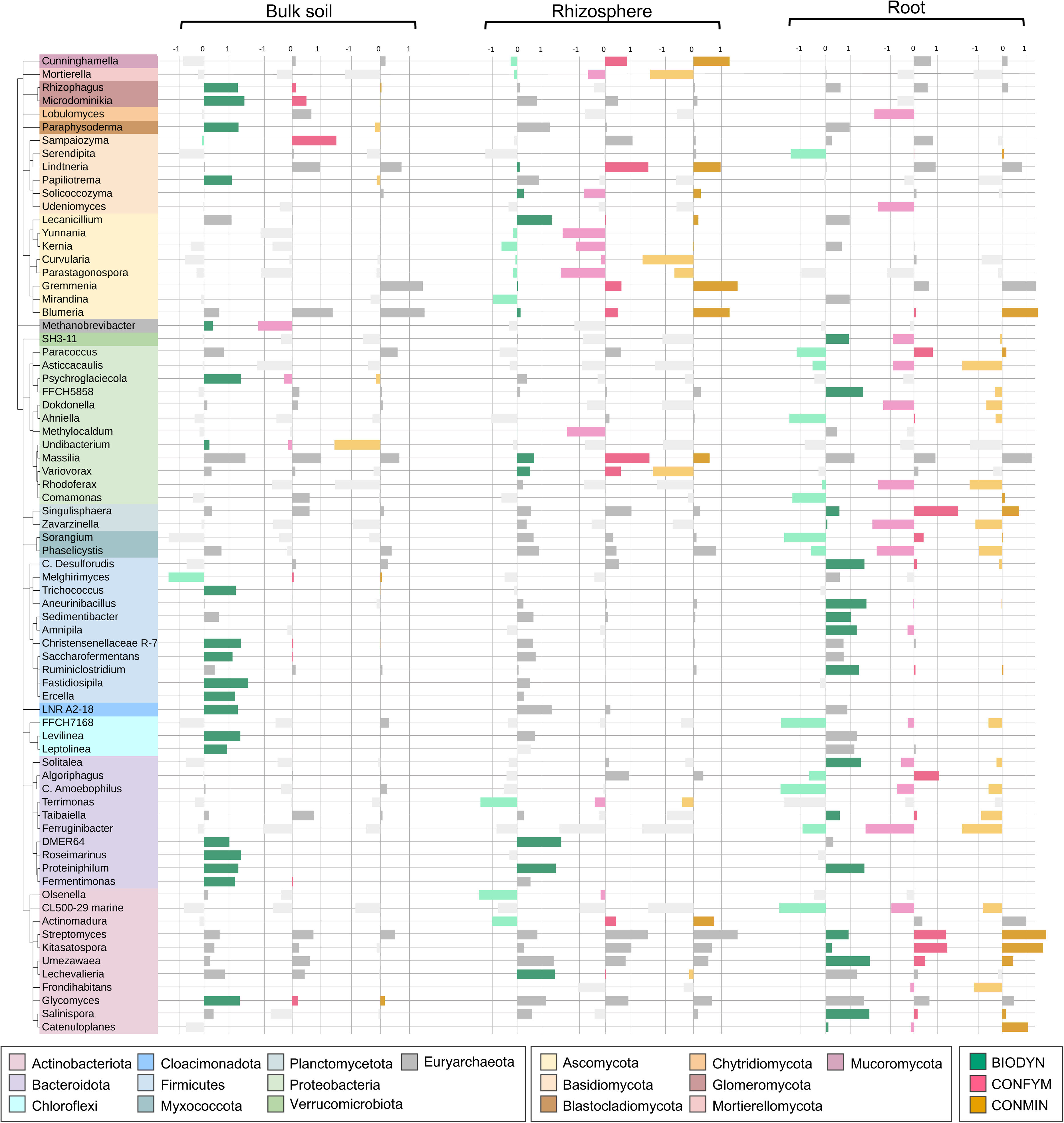
Taxonomic tree displaying prokaryotic and fungal genera in bulk soil, rhizosphere, and roots showing a significant interaction between drought response and cropping system. Genera showing a significant (q < 0.1) interaction are color-coded by the corresponding cropping system, and grey bars are non-significant interactions. Bar plots show the z-transformed relative change in abundance between drought-induced and rainfed treatment of genera enriched or depleted under drought in the respective cropping systems. Color ranges identify corresponding phyla.

Cropping systems had a significant influence on the prokaryotic copiotrophs:oligotrophs ratio in the bulk soil and rhizosphere (Supplementary Figure 12). A significantly higher copiotrophs:oligotrophs ratio was found for drought when compared to the rainfed control in the bulk soil and rhizosphere at the third sampling date. After the rewetting, a higher copiotrophs:oligotrophs ratio was detected (i.e. only measured in bulk soil). A significantly increased ratio of copiotrophs:oligotrophs was found in the roots under drought compared to the control at the second and third sampling date (Supplementary Figure 12).

## Discussion

### Implementation of drought

Drought conditions were successfully induced at field scale (Figure 2, Supplementary Figure 2), with a reduction in water availability characteristic of a comparatively severe drought stress [59,60]. There was no significant effect of cropping system on the decrease in GWC (Figure 2). Although the magnitude of the water content decrease differed between the measurement methods (Figure 2, Supplementary Figure 2), they all showed a continuous decrease in water content in the sheltered plots. A recent short-term, partial sheltering study in two cropping systems of the same field found different GWC reductions between the cropping systems under moderate drought but not under severe drought [59]. Compared with the former study, the drought implemented in the current study was longer, more severe and differences between sheltered and control plots were more pronounced. Studies showed that the effect of SOC content on water retention decreases with decreasing soil water potential [61,62], resulting in little impact on water retention under severe drought. In addition, SOC contents have limited effects on soil water retention in soil rich in silt and clay minerals [61,62]. Since the soil at the DOK trial is a Haplic Luvisol and contains around 72% silt and 16% clay [29], the potential of SOC content to increase the soil water retention in this field experiment is likely limited. Yet, it is important to note that the soil C content in this field experiment is low compared to other agricultural field sites [63], which might further influence the effect of SOC on soil water retention.

The increased temperature of 0.8 ± 0.02 °C below the rainout shelters during winter led to enhanced plant height and biomass at stem elongation. However, drought reduced plant height at flowering and ripening as reported in the literature [64], while dry biomass was not affected and thus contradicting the results of Wittwer et al. (2023) [65]. Khadka et al. (2020) [64] argued that for example, drought-tolerant varieties tend to grow smaller and increase their root biomass to access deeper soil layers. This potentially helped the plants to maintain biomass under drought. At the last sampling date, sheltered wheat plants were overripe potentially resulting in the loss of part of grains before sampling. Nevertheless, there was no significant increase in volunteer grain recorded in fall 2022 in the previously sheltered area (data not shown). It is crucial to mention that plant biomass was measured on a small area (three wheat rows of 17.5 cm × 8 cm), which might not accurately represent yields. Plant height differences between the conventional and biodynamic systems were caused by the application of plant growth regulators in conventional systems. Yet, plants in

BIODYN did not differ in plant height between the water regimes. The grown variety Wiwa was specifically bred for organic cropping systems, which could result in an improved adaptation to organic systems and subsequently better stress tolerance [66]. The impact of drought on plants might depend further on the timing, duration, and severity of the drought, potentially having stronger effects in the early plant stages.

### Drought effect on fungal and prokaryotic communities

Drought altered soil fungal and prokaryotic community structures in all studied compartments although the effect observed in the bulk soil compartment was not very strong (Table 1, Figure 3). Drought effects on microbial communities are in accordance with previous studies reporting on the effects of drought on soil microbes [4,5]. CAP ordinations showed distinct microbial communities between the drought-induced and control plots in all cropping systems (Figure 3), which was largely confirmed by the PERMANOVA results except for prokaryotes in the bulk soil (Table 1); for the latter, effects of drought might have been masked by other more dominant drivers such as cropping system and soil texture. Notably, the effect of drought on microbial communities became stronger in proximity of plant roots, whereas the effect of the cropping system became weaker. Fungal and prokaryotic abundance measured in bulk soil only was not affected by drought (Supplementary Figure 7). Other studies show contrasting results on microbial abundance or biomass [11,12,67,68] ranging from a decrease to no effects or even an increase under drought. The conflicting findings may depend on the evaluation method, soil type, drought severity, and duration. Drought reduced soil respiration in all cropping systems (Supplementary Figure 6). A reduction of soil activity under reduced water availability is commonly reported [11,12]. Interestingly, soil respiration was lowest in the BIODYN treatment. Other studies reported higher respiration rates in organically managed cropping systems, but these studies assessed basal respiration under controlled conditions instead of *in-situ* soil respiration [27,28].

In contrast to our first hypothesis, drought affected soil fungi more strongly than prokaryotes, particularly in bulk soil and rhizosphere (Table 1, Figure 4). Previous studies observed stronger drought effects on prokaryotic community composition [4,5]. Yet, many of these studies were performed either in greenhouse pots or in grasslands, which are managed differently than cropping systems. Fungal hyphal networks are crucial for plant water acquisition [13], and these networks might be more disturbed in cropping systems compared to grasslands by management practices such as soil tillage and mechanical weeding [69]. We did not find a drought effect on fungal abundance, as assessed by rRNA gene copy numbers (Supplementary Figure 7). However, it was shown that hyphal networks do not necessarily contain nucleoid acids and rRNA gene copy numbers might therefore not correlate well with hyphal length [70]. The F/P ratio was lowest in BIODYN in the bulk soil and rhizosphere (Supplementary Figure 7). Since mechanical weeding is performed twice in BIODYN in addition to tillage, the fungal networks might have been disrupted more strongly in this system. Our findings are nevertheless in accordance with a recent spring wheat field experiment, which showed a stronger drought influence on soil fungi compared to prokaryotes [71], argueing that fungi are more sensitive to changes in plant exudation, particularly carbon. Two other field studies in cropping systems with wheat, sugar beet, and maize found a stronger drought response of bacterial communities compared to fungal [4,72], implying that response to drought also depends on other variables such as crop, soil properties, climate, drought severity, and other agricultural practices. Multi-trophic interactions might also influence the microbial drought response such as reduction or shifts of protists or nematodes, which have been shown to be drought-sensitive [73,74]. Such effects might subsequently affect feeding pressure or release nutrients to soils. It is important to mention that a stronger shift of microbial communities in response to drought could also suggest a higher adaptation potential rather than lack of resistance to drought. Another potential explanation for the weaker drought response of prokaryotes compared to fungi could be attributed to preceding summer droughts in 2018 and 2019, which might have led to an adaptation of bacteria to drought, as the fast adaptation of bacteria towards stress is well-known [75]. Prokaryotes might be protected from drought within microaggregates [15], resile in small pores, or become dormant [10,14].

Higher copiotrophic/oligotrophic ratios under drought are contradictory to the hypothesis of Naylor et al. (2018) [16] and previous results in forests and grasslands showing that oligotrophs thrive under drought conditions [76,77]. Opposed to forest and extensively used grassland soils, agricultural cropping systems are frequently fertilized, which might influence how oligotrophs and copiotrophs respond to drought.

In conclusion, this field experiment showed that soil fungi might be more affected by drought in cropping systems compared to prokaryotes possibly due to soil disturbance. It is important to note that microbial drought response further depends on other factors like soil type, texture, aggregation, climate, drought severity, and multi-trophic interactions [5,78].

### Drought effect on microbial communities within compartments

There was a stronger influence of drought on microbial communities more closely associated with plant roots (Table 1, Figure 3), revealing more taxa sensitive to drought in the rhizosphere and root when compared to the bulk soil (Figure 5, Supplementary Data 1). This finding is in accordance with our second hypothesis (ii) and previous studies [4,22,78]. This effect was stronger for prokaryotes than for fungi. The stronger response of root-associated microbiomes is likely caused by a combination of direct effects of water scarcity on the microbes and indirect effects mediated through the drought-affected plants [79]. On the one hand, drought-stressed plants can alter rooting depth and density [80], consequently changing the microbial habitat. On the other hand, metabolic changes in drought-stressed plants can alter rhizodeposition and thereby affect soil microbial communities, especially in proximity of roots [24]. Through this process, plants can select for root microbes that increase plant drought tolerance [24,79]. Moreover, plants accumulate osmolytes in roots to sustain root growth under low soil water potential [81], which might additionally influence root endophytes. However, specific interactions and plant-microbial pathways under drought are still largely unknown, especially under field conditions.

### Cropping-system dependent resistance to drought

Overall, the effects of drought on microbial abundance and community structure were largely independent of the cropping systems, except for the root prokaryotes (Table 1, Supplementary Figure 7), not providing strong support for the third hypothesis (iii). These results contradict previous results, which showed higher bacterial abundance (measured as phospholipid-derived fatty acids) under drought with the addition of composted manure or green waste compared to the control without organic fertilizer [82,83]. Breitkreuz et al. (2021) [78] found significant cropping system effects on the drought response of bacterial composition in a pot experiment using soils from a conventionally and organically managed field trial. The interactive effect was stronger in sandy soils compared to loamy soils [78]; however, no organic fertilizers were applied in the organic cropping system. On the other hand, other studies found no effect of organic management or reduced tillage on the reduction of decomposition activity under drought in field experiments [60,84], which are not necessarily linked to the community structure. A partial, short-term sheltering experiment in the same long-term trial found no strong interactive effect of cropping system and experimental drought under moderate drought [85], supporting our findings. Pot experiments found a few interactions between drought and the addition of organic amendments assessing enzyme activities and microbial composition through phospholipid acids [82,83,86], mentioning a slower drying in amended soils but when reaching the dry state they exhibited similar behaviors.

Although the cropping-system dependent effects of drought on the microbial community were relatively small, several genera showed a system-specific response (Figure 6). *Streptomyces* and *Kitasatospora* were enriched in CONFYM and especially CONMIN under drought compared to BIODYN. Both are potential plant growth promoting (PGP) bacterial genera known to produce the phytohormone auxin, siderophores, and 1-aminocyclopropane-1-carboxylate (ACC) deaminase [87]. Auxin can increase the growth of lateral roots and root hairs [88]. Plant ethylene contents, which can decrease plant and root growth under stress, are reduced by the ACC deaminase and thereby increase tolerance to stress [89]. Siderophores produced by PGP bacteria can solubilize and sequester iron in soils helping plants with the iron uptake and can be involved in the suppression of plant pathogens [90]. *Streptomyces,* often enriched under drought (Figure 5), are considered to be important for plant drought tolerance and are successful in colonizing root tissue under stress [91]. *Actinomadura* known for siderophore and auxin production was additionally enriched in CONFYM and CONMIN compared to BIODYN [87]. Other potential PGP bacteria particularly enriched under drought in CONFYM were *Massilia* and *Paracoccus* [87,92]. *Variovorax*, which was enriched in CONFYM and BIODYN, has been described to improve plant drought tolerance exhibiting similar mechanisms as mentioned above [93]. In the BIODYN treatment, the genera *Aneurinibacillus, Glycomyces, Lechevalieria, Salinispora,* and *Umezawaea* were enriched under drought, which contain species potentially promoting plant growth and are often found in compost [87,94]. Some species in these genera are known for auxin and siderophore production, and ACC deaminase activity [95] but also feature biocontrol activity [94,96,97]. For soil fungi, the genera *Blumeria*, and *Gremmenia* were increased particularly in CONMIN under drought compared to the other cropping systems (Figure 6). Both are potential plant pathogens, and *Blumeria graminis* is known to infest wheat [98,99], indicating that plants in CONMIN under drought might have experienced a higher pathogen pressure. *Lecanicillium*, *Papiliotrema*, *Microdominikia*, and *Rhizophagus* were enriched under drought in BIODYN. These genera are known to contain PGP species [100–104]. Rhizophagus, for example, are arbuscular mycorrhizal fungi known to be able to improve plant drought tolerance [103].

Interestingly, several genera that increased under drought in BIODYN compared to the other cropping systems are known to contain facultatively or obligate anaerobic species (i.e. *Fermentimonas, Proteiniphilum, Roseimarinus, Solitalea, Leptolinea, Levilinea, Ercella, Fastidiosipila, Ruminiclostridium, Saccharofermentans, Christensenellaceae R-7 group, Sedimentibacter, Candidatus Desulforudis, Trichococcus*, *Methanobrevibacter;* Figure 6) [105,106]. Many of these genera have been found in slurry or animal rumen and are involved in fermentation and methanogenesis [107,108]. Indeed, slurry was applied in February and March in the BIODYN treatment but not in CONFYM and CONMIN. However, this relative increase of species involved in methanogenesis in BIODYN soils under drought did not increase *in-situ* methane emissions (data not shown), which suggests that the increased relative abundance did not translate into increased activity, either because these genera were inactive or dead [60,109].

In this study, we defined resistance as the ability to tolerate drought by not changing community composition [26]. Hence, a more pronounced shift in microbial community structure upon drought would suggest lower resistance to drought, while no or a small shift would indicate stronger resistance. However, it remains uncertain whether increases or decreases of specific taxa in one versus the other cropping system implies lower resistances in one system than the other, or if it actually represents some adaptation mechanisms.

In conclusion, all cropping systems showed under drought enrichments of some PGP genera potentially involved in the improvement of plant drought tolerance, especially of the phylum *Actinobacteriota*. Generally, fungal genera possibly involved in improving plant drought tolerance were enriched in BIODYN. Moreover, microbial communities were similarly affected by drought in all cropping systems. Hence, we found no clear indication that the application of composted or stacked manure in BIODYN and CONFYM, the associated increase in SOC [29] and microbial diversity [25], the reduction of pesticide application, or other factors like the biodynamic preparations in BIODYN could increase microbial resistance to drought. Additionally, this long-term field trial already includes some regenerative practices such as shallow tillage, cover cropping, and incorporation of grass-clover into the crop rotation in all cropping systems. Those practices might have already improved microbial resistance to drought and still, shifts of microbial communities were recorded. However, we did not find a strong indication of different resistances of the microbial communities, and GWC reduction did not differ under drought between the manure-treated and minerally fertilized systems. Yet, cropping systems still harbour distinct microbiomes under severe drought and these distinct communities might feature contrasting potentials to cope with drought. It is important to note that this study is confined to one climate, crop, and soil type.

### Cropping system-dependent resilience to drought

Despite the effect of drought on the bulk soil microbiome was not very strong, a drought legacy effect one week and about two months after rewetting was clearly detectable (Figure 4, Supplementary Figure 11, Supplementary Table 6), which is supported by previous studies [6,91]. However, prokaryotic and fungal communities did not show distinct resilience patterns depending on the cropping system. Therefore, we have to reject our fourth hypothesis that different cropping systems might show different capacities for resilience (iv). Some pot studies found comparable resilience in soils with and without organic amendments assessed by enzyme activities, basal respiration, and phospholipid acids [83,86], while another study found differences in resilience patterns using molecular analysis [110].

There is a limited number of studies that have assessed microbial resilience to drought in contrasting cropping systems, particularly involving plants and at field-scale. This study indicates that the application of organic amendments in the form of farmyard manure in organic and mixed conventional cropping systems, or the reduction of pesticide application or factors like biodynamic preparations might have limited effects on microbial resilience after drought. This is supported by the finding that we did not find increased soil moisture in one over the other cropping systems after rewetting (Figure 1). However, the effect may depend on climatic conditions, soil type, and crop.

## Conclusions

First, our results suggest that in cropping systems soil fungi might be less resistant to drought compared to prokaryotes possibly because of frequent soil disturbances or stronger interaction with plant exudates. Secondly, this study indicates that cropping systems considered to promote soil biodiversity and SOC content, such as organic cropping systems, might not be able to mitigate the impact of severe drought on soil biodiversity. Hence, alternative farming practices might have to be included to enhance microbial resistance and resilience in cropping systems. Given that this field trial already includes some regenerative practices in all cropping systems, comparison to other cropping systems including more conventional practices such as conventional tillage, fallows, or monocropping would put the cropping systems in the DOK trial into a broader perspective. Since this study focused on assessing the effects of drought on taxonomic diversity, our conclusions about microbiome-mediated changes in soil functions under drought are still limited. This emphasizes to study the effects of drought on soil functions with - for example - metagenomics. Finally, stronger drought effects were found for microbes more closely associated with roots, which emphasizes the importance of plant-microbe interactions. Additional studies are needed to examine rhizodeposition patterns under drought in different cropping systems in order to better understand the relevance of these interactions to mitigate the impact of climate change stressors.

## Data Availability

Raw sequence data were deposited in the European Nucleotide Archive under the accession number PRJEB73799.

## Acknowledgment

We thank members of the research groups at FiBL Frick (Group Soil fertility & Climate), Agroscope Zürich (Group Water Protection and Substance Flows), Uni Kassel (Group soil biology and plant nutrient), and ETH Zürich (Sustainable Agroecosystems Group) for their contributions to this study. We are especially grateful to Hans-Ulrich Zbinden, Frédéric Perrochet, Moritz Sauter, Adrian Lustenberger, Matti Barthel, Charles Nwokoro, Tim Juchli, Noah Schweizer, Moritz Bach, and Bernhard Stehle for their support during fieldwork. We are grateful to all the helpers during sampling and *in-situ* data collection including Matthias Lang, Sabrina Niehaus, Juliana Jäggle, Sarah Symanczik, Marijn Van de Broek, Lian Tengxiang, Tania Galindo, Tabata Bublitz, and Astrid Jäger. We are also grateful to Rafaela Feola Conz, Matti Barthel, Britta Jahn-Humphrey, and the soil and elemental analysis group at Agroscope for their technical and work support in the laboratory. Finally, we would like to acknowledge Maria Domenica Moccia at the Functional Genomics Center Zurich (FGCZ) for the amplicon sequencing service on the Illumina MiSeq v3 platform.

## Authors Contributions

MH, JM, and PM designed the field experiment. EK and DK managed the sheltering experiment and sampling. EK and RFC carried out the molecular lab work. EK performed the bioinformatics, statistical analysis, and data visualization. EK and MH wrote and revised the original draft, all authors edited the manuscript. MH, JM, and JS supervised the research.

