## Supplementary Material for "Microbial resistance and resilience to drought under organic and conventional farming"

### Supplementary Information

Supplementary Table 1: Detailed timeline of the agricultural interventions in rainout-sheltering experiment in the DOK long-term field trial. Samples were collected at five timepoints (in bold), three during the wheat vegetation period/drought phase and two after the vegetation period/rewetting.

| 7 October 2021 | Composted manure application in BIODYN (10 t ha^-1^)  Stacked manure application in CONFYM (12 t ha^-1^)  Mineral fertilizer application in CONMIN (29 kg P ha ^-1^, 75 kg K ha^-1^) |
| --- | --- |
| 15 October 2021 | Primary tillage with disk harrow in all systems |
| 19 October 2021 | Seedbed preparation with rotary harrow and sowing of winter wheat variety Wiwa in all systems (dressed seed in conventionally managed systems) |
| 15 – 17 November 2021 | Installation of rainout-shelter with foil |
| 1 – 15 December 2021 | Installation TDR, TOMST, PAR sensors  Installation of rain gutters and IBC tanks |
| 22 December 2021 | Connection lower side of rain gutters with the pipe system |
| 6 January 2022 | Installation of respiration chambers |
| 20 January 2022 | Connection higher side of rain gutters to IBC tanks and pipe system |
| 28 January –  2 February 2022 | Irrigation sheltered plots with 30 mm |
| 18 February 2022 | Irrigation sheltered plots with 15 mm |
| 1 March 2022 | Irrigation sheltered plots with 10 mm |
| 10 March 2022 | Mechanical weeding by hand in the BIODYN |
| 15 March 2022 | Biodynamic preparations in BIODYN (Horn manure 96 g ha^.1^) |
| 16 March 2022 | First slurry application in BIODYN (2.5 l ha^-1^)  First mineral N application in conventionally managed systems (60 kg N ha^-1^, 14 kg P ha^.1^, 14 kg K ha^.1^ in CONFYM and 70 kg N ha^-1^ in CONMIN) |
| 28 March 2022 | Application of herbicide (Othello ® Star; Bayer AG, Leverkusen, Deutschland) and plant growth regulators (CCC 720 ®; Bayer AG) in conventionally managed systems (1 and 0.5 l ha^-1^, respectively) |
| 31 March 2022 | Irrigation sheltered plots with 2 mm  Installation of HOBO sensors |
| 5 April 2022 | CO_2_, N_2_O, CH_4_ measurement |
| 11 April 2022 | Application of plant-growth regulator (Moddus; Syngenta, Basel, Switzerland) in conventionally managed systems (0.6 l ha^-1^) |
| 12 April 2022 | CO_2_, N_2_O, CH_4_ measurement |
| 13 April 2022 | Second slurry application in BIODYN (4 l ha^-1^)  Second mineral N application in conventionally managed systems (30 kg N ha^-1^ in CONFYM and 40 kg N ha^-1^ in CONMIN) |
| 19 April 2022 | CO_2_, N_2_O, CH_4_ measurement |
| 21 April 2022 | Mechanical weeding by hand in BIODYN |
| 26 April 2022 | CO_2_, N_2_O, CH_4_ measurement |
| **27 – 28 April 2022** | **First sampling starting stem elongation** |
| 3 May 2022 | CO_2_, N_2_O, CH_4_ measurement |
| 5 May 2022 | Biodynamic preparations in BIODYN (Horn silica 8 g ha^-1^) |
| 10 May 2022 | CO_2_, N_2_O, CH_4_ measurement |
| 17 May 2022 | CO_2_, N_2_O, CH_4_ measurement  Third mineral N application in conventionally managed systems (40 kg N ha^-1^ in CONFYM and 40 kg N ha^-1^ in CONMIN) |
| 18 May 2022 | Application of fungicide (Aviator® Xproin; Bayer AG) conventionally managed systems (1.25 l ha^-1^) |
| 24 May 2022 | CO_2_, N_2_O, CH_4_ measurement |
| 28 May 2022 | Biodynamic preparations in BIODYN (Horn silica 8 g ha^-1^) |
| 31 May 2022 | CO_2_, N_2_O, CH_4_ measurement |
| **31 May – 1 June 2022** | **Second sampling at flowering** |
| 10 June 2022 | CO_2_, N_2_O, CH_4_ measurement |
| 14 June 2022 | CO_2_, N_2_O, CH_4_ measurement |
| 17 June 2022 | Distribution of net against bird damage |
| 21 June 2022 | CO_2_, N_2_O, CH_4_ measurement |
| 28 June 2022 | CO_2_, N_2_O, CH_4_ measurement |
| **5 July 2022** | **Third sampling at ripening**  CO_2_, N_2_O, CH_4_ measurement |
| 6 – 7 July 2022 | Removing rainout-shelters and HOBO, TDR, PAR sensors |
| 13 July 2022 | Harvest of total plot |
| 14 July 2022 | Rewetting with 36 mm tap water |
| 15 July 2022 | CO_2_, N_2_O, CH_4_ measurement |
| 19 July 2022 | Straw harvest |
| **20 July 2022** | **Fourth sampling one week after rewetting**  CO_2_, N_2_O, CH_4_ measurement  Removal of TOMST sensors |
| 21 July 2022 | Mulching stubbles |
| 22 July 2022 | Stubble processing with rotary harrow |
| 27 July 2022 | Composted manure in BIODYN (5 t ha^-1^) |
| 28 July 2022 | Manual weeding with cultivator in all systems |
| 29 July 2022 | Mineral N application in conventionally managed systems (30 kg N ha^-1^ in CONFYM  and 30 kg N ha^-1^ in CONMIN) |
| 16 August 2022 | Seedbed preparation with disk harrow in all systems and sowing of green manure in all  systems |
| 20 August 2022 | Biodynamic preparations in BIODYN (Horn silica 8 g ha^-1^) |
| **13 September 2022** | **Fifth sampling eleven weeks after rewetting** |

Supplementary Table 2: Summary of all PCR and qPCR conditions including primer sequences, cycling conditions, PCR reagents, and reference.

| Analysis | Primer | Cycling | | | | Mastermix | Ref |
| --- | --- | --- | --- | --- | --- | --- | --- |
| PCR 16S in bulk soil, rhizosphere | 341F (5’-CCTAYGGGDBGCWSCAG-3’)  806R (5’-GGACTACNVGGGTHTCTAAT-3’) | 1 | 2 min | 95 °C |  | 1x GoTaq® G2 Hot start Master mix  0.5 mM MgCl_2_  0.4 µM primer F  0.4 µM primer R  40 ng DNA | (1) |
|  |  | 2 | 40 sec | 95 °C |  |  |  |
|  |  | 3 | 40 sec | 58 °C |  |  |  |
|  |  | 4 | 1 min | 72 °C | 30 × |  |  |
|  |  | 5 | 10 min | 72 °C |  |  |  |
| PCR ITS in bulk soil, rhizosphere, root | 5.85-Fung (5'-AACTTTYRRCAAYGGATCWCT-3′)  ITS4-Fung (5'-AGCCTCCGCTTATTGATATGCTTAART-3′) | 1 | 2 min | 95 °C |  | 1x GoTaq® G2 Hot start Master mix  1 mM MgCl_2_  0.4 µM primer F  0.4 µM primer R  40 ng DNA | (2) |
|  |  | 2 | 40 sec | 95 °C |  |  |  |
|  |  | 3 | 40 sec | 58 °C |  |  |  |
|  |  | 4 | 1 min | 72 °C | 35 × |  |  |
|  |  | 5 | 10 min | 72 °C |  |  |  |
| PCR 16S root | 341F (5’-CCTAYGGGDBGCWSCAG-3’)  806R (5’-GGACTACNVGGGTHTCTAAT-3’)  mPNA (5’-GGCAAGTGTTCTTCGGA-3’)  pPNA (5’-GGCTCAACCCTGGACAG-3’) | 1 | 3 min | 95 °C |  | 1x GoTaq® G2 Hot start Master mix  0.5 mM MgCl2  0.4 µM primer F  0.4 µM primer R  0.76 µM mPNA  0.76 µM pPNA  40 ng DNA | (1,3) |
|  |  | 2 | 40 sec | 95 °C |  |  |  |
|  |  | 3 | 10 sec | 78 °C |  |  |  |
|  |  | 4 | 40 sec | 58 °C |  |  |  |
|  |  | 5 | 1 min | 72 °C | 35 × |  |  |
|  |  | 6 | 10 min | 72 °C |  |  |  |
| qPCR inhibition test | SP6 (5'-ATTTAGGTGACACTATAG-3')  T7 (5'-TAATACGACTCACTATAGGG-3') | 1 | 3 min | 98 °C |  | 1x SsoAdvanced universal SYBR® Green supermix  0.75 µM primer F  0.75 µM primer R  20 ng DNA  10^4^ copy number pGEM-T plasmid | (4) |
|  |  | 2 | 15 sec | 95 °C |  |  |  |
|  |  | 3 | 30 sec | 48 °C |  |  |  |
|  |  | 4 | 30 sec | 72 °C |  |  |  |
|  |  | 5 | 15 sec | 80 °C | 30 × |  |  |
|  |  | 6 | 5 sec | 75-95 °C | 0.3 °C |  |  |
| qPCR 16S in bulk soil, rhizosphere | 515Y-F (modified) (5'-GTGYCAGCMGCCGCGGTAA-3')  806R (original) (5'-GGACTACHVGGGTWTCTAAT-3') | 1 | 3 min | 98 °C |  | 1x SsoAdvanced universal SYBR® Green supermix  0.75 µM primer F  0.75 µM primer R  20 ng DNA | (1,5) |
|  |  | 2 | 15 sec | 95 °C |  |  |  |
|  |  | 3 | 30 sec | 52 °C |  |  |  |
|  |  | 4 | 30 sec | 72 °C |  |  |  |
|  |  | 5 | 15 sec | 80 °C | 35 × |  |  |
|  |  | 6 | 5 sec | 75-95 °C | 0.3 °C |  |  |
| qPCR 18S in bulk soil, rhizosphere | FR1 (5'-ANCCATTCAATCGGTANT-3')  FF390 (5'-CGATAACGAACGAGACC-3') | 1 | 3 min | 98 °C |  |  | (6) |
|  |  | 2 | 20 sec | 95 °C |  |  |  |
|  |  | 3 | 30 sec | 51 °C |  |  |  |
|  |  | 4 | 30 sec | 72 °C |  |  |  |
|  |  | 5 | 15 sec | 80 °C | 35 × |  |  |
|  |  | 6 | 5 sec | 75-95 °C | 0.3 °C |  |  |

Supplementary Table 3: A) Effect of water regime, cropping system, and sampling dates on the total soil carbon, total soil nitrogen, plant-available phosphorus, plant-available potassium, plant-available magnesium, and pH assessed by PERMANOVA based on Euclidian distances (F-ratio, p-value, and R^2^) in the bulk soil **during the drought period**. Values p < 0.05 are indicated as bold. Heteroscedasticities are indicated as superscript ^1^. B) Average values (mean ± se) for each cropping system and water regime. Different letters represent significant differences at p < 0.05 corrected for multiple comparisons using the Benjamini–Hochberg method.

| PERMANOVA^A^ | | Total soil carbon | | | | Total soil nitrogen | | | | Plant-available phosphorus | | | | Plant-available potassium | | | | Plant-available magnesium | | | | pH | | |
| --- | --- | --- | --- | --- | --- | --- | --- | --- | --- | --- | --- | --- | --- | --- | --- | --- | --- | --- | --- | --- | --- | --- | --- | --- |
|  | F (p) | | | R^2^ | F (p) | | | R^2^ | F (p) | | | R^2^ | F (p) | | | R^2^ | F (p) | | | R^2^ | F (p) | | | R^2^ |
| Water regime (W) | 0.63 (0.4295) | | | 0.005 | 0.00 (0.9652) | | | 0.000 | 6.51 (**0.0129**) | | | 0.059 | 18.24 (**0.0001**) | | | 0.185 | 0.90 (0.3473) | | | 0.006 | 0.08 (0.7821) | | | 0.000 |
| Cropping System (C) | 30.06 (**0.0001**)^1^ | | | 0.488 | 5.78 (**0.0013**) | | | 0.155 | 18.34 (**0.0001**) | | | 0.332 | 6.22 (**0.0023**) | | | 0.126 | 45.68 (**0.0001**)^1^ | | | 0.594 | 120.55 (**0.0001**)^1^ | | | 0.786 |
| Sampling Date (S) | 0.05 (0.8231) | | | 0.000 | 0.91 (0.3610) | | | 0.012 | 2.60 (0.1055) | | | 0.024 | 0.08 (0.7718) | | | 0.001 | 0.67 (0.4210) | | | 0.004 | 0.53 (0.4667) | | | 0.002 |
| W x C | 1.04 (0.3537) | | | 0.017 | 0.96 (0.4160) | | | 0.026 | 0.28 (0.7664) | | | 0.005 | 3.22 (**0.0458)** | | | 0.065 | 0.16 (0.8532) | | | 0.002 | 1.99 (0.1414) | | | 0.013 |
| W x S | 0.04 (0.8334) | | | 0.000 | 0.01 (0.9395) | | | 0.000 | 2.07 (0.1542) | | | 0.019 | 0.13 (0.7202) | | | 0.001 | 0.00 (0.9724) | | | 0.000 | 0.39 (0.5354) | | | 0.001 |
| C x S | 0.04 (0.9584) | | | 0.001 | 0.00 (0.9978) | | | 0.000 | 1.06 (0.3557) | | | 0.019 | 0.43 (0.6602) | | | 0.009 | 0.09 (0.9081) | | | 0.001 | 0.31 (0.7351) | | | 0.002 |
| W x C x S | 0.07 (0.9353) | | | 0.001 | 0.01 (0.9900) | | | 0.000 | 0.01 (0.9893) | | | 0.000 | 0.32 (0.7275) | | | 0.006 | 0.16 (0.8538) | | | 0.002 | 0.09 (0.9125) | | | 0.001 |
| AVERAGE^B^ | | | Total soil carbon  (mg/kg) | | | | Total soil nitrogen (mg/kg) | | | | Plant-available phosphorus (mg/kg) | | | | Plant-available potassium (mg/kg) | | | | Plant-available magnesium (mg/kg) | | | | pH | |
|  | | | Mean ± se | | | | Mean ± se | | | | Mean ± se | | | | Mean ± se | | | | Mean ± se | | | | Mean ± se | |
| BIODYN control | | | 1.70 ± 0.03 a | | | | 0.186 ± 0.004 a | | | | 1.04 ± 0.10 ab | | | | 10.7 ± 0.9 abc | | | | 142 ± 5 a | | | | 6.81 ± 0.03 a | |
| CONFYM control | | | 1.45 ± 0.02 b | | | | 0.177 ± 0.020 ab | | | | 1.41 ± 0.08 cd | | | | 8.5 ± 0.5 ad | | | | 113 ± 2 b | | | | 6.09 ± 0.05 b | |
| CONMIN control | | | 1.42 ± 0.06 b | | | | 0.156 ± 0.008 b | | | | 0.87 ± 0.04 a | | | | 6.9 ± 0.6 d | | | | 159 ± 6 ac | | | | 6.13 ± 0.08 b | |
| BIODYN drought-induced | | | 1.82 ± 0.06 a | | | | 0.199 ± 0.007 a | | | | 1.16 ± 0.05 b | | | | 11.4 ± 0.7 bc | | | | 143 ± 6 a | | | | 6.76 ± 0.02 a | |
| CONFYM drought-induced | | | 1.45 ± 0.02 b | | | | 0.161 ± 0.003 b | | | | 1.60 ± 0.13 c | | | | 13.6 ± 1.1 b | | | | 119 ± 2 b | | | | 6.19 ± 0.03 b | |
| COMIN drought-induced | | | 1.40 ± 0.07 b | | | | 0.160 ± 0.009 b | | | | 1.12 ± 0.10 abd | | | | 10.0 ± 0.9 ac | | | | 163 ± 5 c | | | | 6.04 ± 0.06 b | |

Supplementary Table 4: A) Effect of water regime, cropping system, and sampling dates on the total soil carbon, total soil nitrogen, plant-available phosphorus, plant-available potassium, plant-available magnesium, and pH assessed by PERMANOVA based on Euclidian distances (F-ratio, p-value, and R^2^) in the bulk soil **after rewetting**. Values p < 0.05 are indicated as bold. Heteroscedasticities are indicated as superscript ^1^. B) Average values (mean ± se) for each cropping system and water regime. Different letters represent significant differences at p < 0.05 corrected for multiple comparisons using the Benjamini–Hochberg method.

| PERMANOVA^A^ | | Total soil carbon | | | | Total soil nitrogen | | | | Plant-available phosphorus | | | | Plant-available potassium | | | | Plant-available magnesium | | | | pH | | |
| --- | --- | --- | --- | --- | --- | --- | --- | --- | --- | --- | --- | --- | --- | --- | --- | --- | --- | --- | --- | --- | --- | --- | --- | --- |
|  | F (p) | | | R^2^ | F (p) | | | R^2^ | F (p) | | | R^2^ | F (p) | | | R^2^ | F (p) | | | R^2^ | F (p) | | | R^2^ |
| Water regime (W) | 0.39 (0.5342) | | | 0.005 | 0.85 (0.3712) | | | 0.013 | 0.47 (0.5057) | | | 0.007 | 2.60 (0.1176) | | | 0.041 | 0.17 (0.6762) | | | 0.002 | 0.92 (0.3375) | | | 0.005 |
| Cropping System (C) | 20.51 (**0.0001**)^1^ | | | 0.521 | 15.02 (**0.0003**)^1^ | | | 0.443 | 9.44 (**0.0004**) | | | 0.285 | 7.93 (**0.0011**) | | | 0.251 | 27.06 (**0.0001**)^1^ | | | 0.595 | 70.45 (**0.0001**)^1^ | | | 0.770 |
| Sampling Date (S) | 0.20 (0.6614) | | | 0.003 | 0.03 (0.8617) | | | 0.000 | 4.28 (**0.0466)** | | | 0.065 | 0.01 (0.9341) | | | 0.000 | 0.09 (0.7639) | | | 0.001 | 2.74 (0.1050) | | | 0.015 |
| W x C | 0.49 (0.6168) | | | 0.012 | 0.21 (0.8108) | | | 0.006 | 1.29 (0.2877) | | | 0.039 | 0.24 (0.7865) | | | 0.007 | 0.07 (0.9326) | | | 0.002 | 0.79 (0.4506) | | | 0.009 |
| W x S | 0.06 (0.8040) | | | 0.001 | 0.19 (0.6600) | | | 0.003 | 0.00 (1.0000) | | | 0.000 | 4.11 (0.0508) | | | 0.065 | 0.11 (0.7428) | | | 0.001 | 0.11 (0.7349)) | | | 0.004 |
| C x S | 0.03 (0.9686) | | | 0.001 | 0.07 (0.9358) | | | 0.002 | 1.20 (0.3047) | | | 0.036 | 0.52 (0.5968) | | | 0.017 | 0.02 (0.9832) | | | 0.000 | 0.38 (0.6830) | | | 0.004 |
| W x C x S | 0.02 (0.9794) | | | 0.001 | 0.07 (0.9351) | | | 0.002 | 0.85 (0.4380) | | | 0.026 | 1.51 (0.2271) | | | 0.048 | 0.17 (0.8440) | | | 0.004 | 0.02 (0.9843) | | | 0.000 |
| AVERAGE^B^ | | | Total soil carbon  (mg/kg) | | | | Total soil nitrogen (mg/kg) | | | | Plant-available phosphorus (mg/kg) | | | | Plant-available potassium (mg/kg) | | | | Plant-available magnesium (mg/kg) | | | | pH | |
|  | | | Mean ± se | | | | Mean ± se | | | | Mean ± se | | | | Mean ± se | | | | Mean ± se | | | | Mean ± se | |
| BIODYN control | | | 1.75 ± 0.05 a | | | | 0.196 ± 0.006 ab | | | | 1.38 ± 0.11 a | | | | 14.2 ± 1.4 a | | | | 143 ± 5 a | | | | 6.84 ± 0.03 a | |
| CONFYM control | | | 1.45 ± 0.02 b | | | | 0.163 ± 0.001 c | | | | 1.37 ± 0.10 a | | | | 11.2 ± 1.3 ab | | | | 116 ± 2 b | | | | 6.20 ± 0.04 b | |
| CONMIN control | | | 1.40 ± 0.08 b | | | | 0.160 ± 0.010 c | | | | 0.85 ± 0.08 b | | | | 8.3 ± 1.1 b | | | | 163 ± 8 a | | | | 6.18 ± 0.12 b | |
| BIODYN drought-induced | | | 1.86 ± 0.08 a | | | | 0.208 ± 0.009 a | | | | 1.44 ± 0.15 a | | | | 15.7 ± 1.7 a | | | | 148 ± 7 a | | | | 6.88 ± 0.03 a | |
| CONFYM drought-induced | | | 1.42 ± 0.03 b | | | | 0.164 ± 0.003 c | | | | 1.26 ± 0.09 a | | | | 12.1 ± 0.6 a | | | | 119 ± 2 b | | | | 6.14 ± 0.05 b | |
| COMIN drought-induced | | | 1.42 ± 0.09 b | | | | 0.165 ± 0.011 bc | | | | 1.08 ± 0.11 ab | | | | 11.1 ± 1.7 ab | | | | 163 ± 7 a | | | | 6.03 ± 0.09 b | |

Supplementary Table 5: A) Effect of water regime, cropping system, and sampling dates on the prokaryotic and fungal α-diversity (i.e. Shannon index based on iteratively rarefied data) assessed by PERMANOVA based on Euclidean distances (F-ratio, p-value, and R2) in the three compartments (i.e. bulk soil, rhizosphere, and root). Values p < 0.05 are indicated as bold. B) Average Shannon index values (mean ± se) for each cropping system and water regime.

| PERMANOVA^A^ | | Prokaryotes | | | | | | | | | | | Fungi | | | | | |
| --- | --- | --- | --- | --- | --- | --- | --- | --- | --- | --- | --- | --- | --- | --- | --- | --- | --- | --- |
|  | | Bulk soil | | | | Rhizosphere | | | | Root | | | Bulk soil | | Rhizosphere | | Root | |
|  | F (p) | | | R^2^ | F (p) | | | R^2^ | F (p) | | | R^2^ | F (p) | R^2^ | F (p) | R^2^ | F (p) | R^2^ |
| Water regime (W) | 0.07 (0.8042) | | | 0.001 | 2.04 (0.1563) | | | 0.030 | 0.90 (0.3387) | | | 0.008 | 1.58 (0.2091) | 0.016 | 7.65 (**0.0050**) | 0.094 | 15.22 (**0.0004**) | 0.157 |
| Cropping System (C) | 2.69 (0.0770) | | | 0.078 | 2.97 (0.0601) | | | 0.086 | 10.35 (**0.0005**) | | | 0.173 | 12.28 (**0.0001**) | 0.255 | 4.94 (**0.0086**) | 0.121 | 1.39 (0.2596) | 0.029 |
| Sampling Date (S) | 2.48 (0.1208) | | | 0.036 | 0.00 (0.9636) | | | 0.002 | 31.98 (**0.0001**) | | | 0.267 | 1.44 (0.2297) | 0.015 | 1.79 (0.1777) | 0.044 | 6.27 (**0.0037**) | 0.129 |
| W x C | 0.06 (0.9395) | | | 0.002 | 0.22 (0.8097) | | | 0.216 | 1.24 (0.2916) | | | 0.021 | 0.10 (0.9032) | 0.002 | 0.97 (0.3972) | 0.024 | 1.36 (0.2715) | 0.028 |
| W x S | 0.02 (0.8807) | | | 0.000 | 0.30 (0.5924) | | | 0.302 | 1.13 (0.2972) | | | 0.009 | 3.55 (0.0624) | 0.037 | 0.32 (0.7358) | 0.008 | 1.31 (0.2803) | 0.027 |
| C x S | 0.29 (0.7507) | | | 0.008 | 0.62 (0.5346) | | | 0.616 | 0.31 (0.7325) | | | 0.005 | 1.66 (0.1957) | 0.035 | 0.43 (0.7949) | 0.021 | 1.08 (0.3700) | 0.045 |
| W x C x S | 0.01 (0.9950) | | | 0.000 | 0.42 (0.6619) | | | 0.415 | 0.89 (0.4159) | | | 0.015 | 1.28 (0.2882) | 0.027 | 0.49 (0.7580) | 0.024 | 0.67 (0.6230) | 0.028 |
| AVERAGE^B^ | | | Prokaryotes | | | | | | | | | | Fungi | | | | | |
|  | | | Bulk soil | | | | Rhizosphere | | | | Root | | Bulk soil | | Rhizosphere | | Root | |
|  | | | Mean ± se | | | | Mean ± se | | | | Mean ± se | | Mean ± se | | Mean ± se | | Mean ± se | |
| BIODYN control | | | 8.22 ± 0.01 | | | | 8.21 ± 0.01 | | | | 6.98 ± 0.07 | | 4.90 ± 0.07 | | 4.93 ± 0.05 | | 3.47 ± 0.09 | |
| CONFYM control | | | 8.18 ± 0.02 | | | | 8.18 ± 0.02 | | | | 6.77 ± 0.12 | | 4.82 ± 0.03 | | 4.80 ± 0.02 | | 3.63 ± 0.10 | |
| CONMIN control | | | 8.14 ± 0.03 | | | | 8.15 ± 0.03 | | | | 6.69 ± 0.03 | | 4.77 ± 0.04 | | 4.80 ± 0.02 | | 3.71 ± 0.07 | |
| BIODYN drought-induced | | | 8.19 ± 0.02 | | | | 8.17 ± 0.03 | | | | 7.02 ± 0.08 | | 4.88 ± 0.05 | | 4.88 ± 0.05 | | 3.85 ± 0.08 | |
| CONFYM drought-induced | | | 8.13 ± 0.03 | | | | 8.17 ± 0.02 | | | | 6.95 ± 0.07 | | 4.75 ± 0.03 | | 4.72± 0.07 | | 3.80 ± 0.07 | |
| COMIN drought-induced | | | 8.10 ± 0.03 | | | | 8.10 ± 0.04 | | | | 6.64 ± 0.09 | | 4.61 ± 0.09 | | 4.61 ± 0.11 | | 3.87 ± 0.07 | |

Supplementary Table 6: PERMANOVA results (F-ratio, p-value, and R^2^) showing the effect of water regime, cropping system, and sampling dates on the prokaryotic and fungal β-diversity based on Bray-Curtis dissimilarity in the bulk soil after rewetting. Heteroscedasticities are indicated as superscript ^1^. Values p < 0.05 are indicated as bold.

| PERMANOVA | Prokaryotes | | Fungi | |
| --- | --- | --- | --- | --- |
|  | F (p) | R^2^ | F (p) | R^2^ |
| Water regime (W) | 1.9 (0.0545) | 0.030 | 3.7 (**0.0001**) | 0.054 |
| Cropping System (C) | 10.6 (**0.0001**)^1^ | 0.329 | 7.0 (**0.0001**)^1^ | 0.207 |
| Sampling Date (S) | 1.2 (0.1923) | 0.019 | 6.4 (**0.0001)** | 0.094 |
| W x C | 0.9 (0.5659) | 0.027 | 1.1 (0.2580) | 0.033 |
| W x S | 0.8 (0.6396) | 0.012 | 2.0 (**0.0054**) | 0.029 |
| C x S | 0.7 (0.8130) | 0.022 | 1.0 (0.3681) | 0.031 |
| W x C x S | 0.6 (0.9325) | 0.020 | 0.8 (0.9004) | 0.023 |

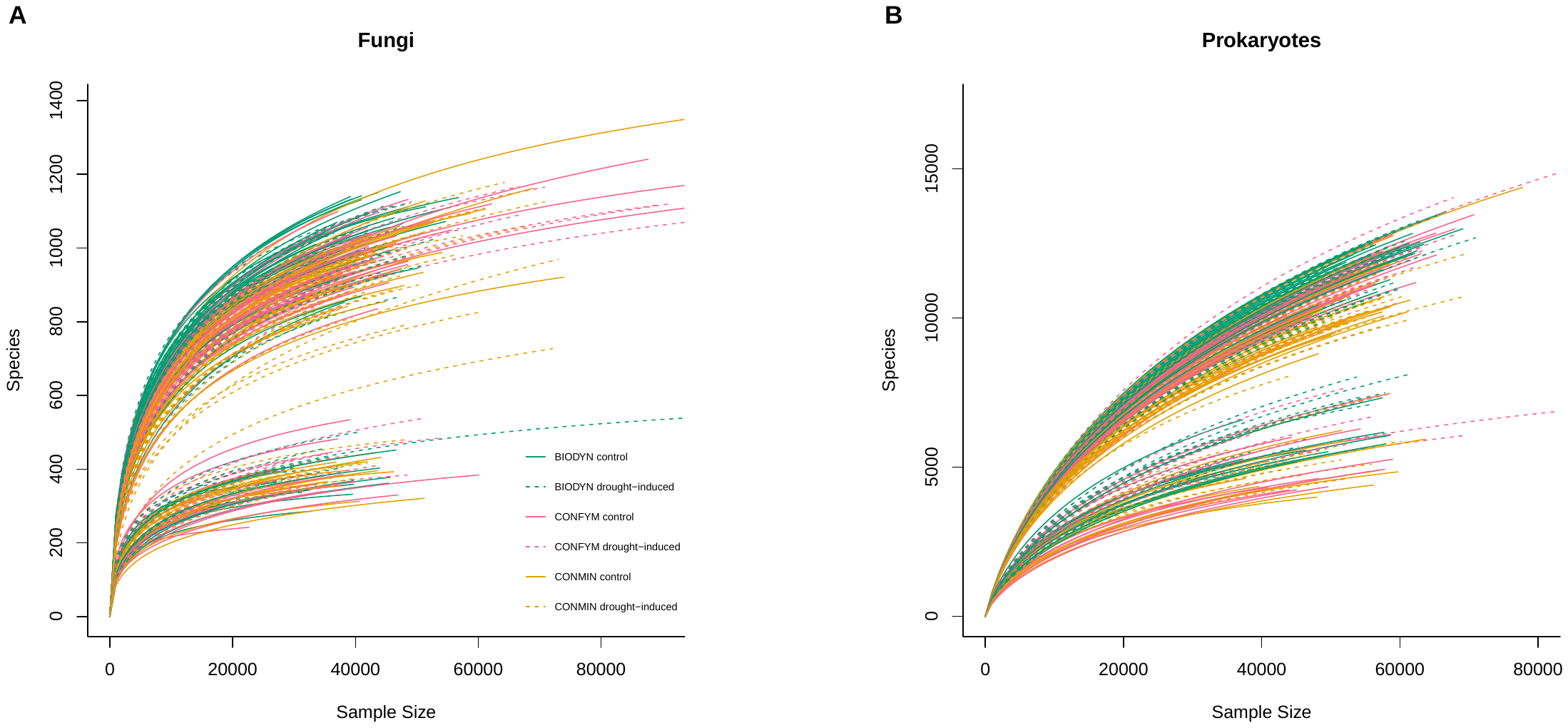

Supplementary Figure 1: Rarefaction curves for all 263 fungal (A) and 261 prokaryotic (B) samples. Samples with very low read counts were excluded (one fungal and three prokaryotic samples).

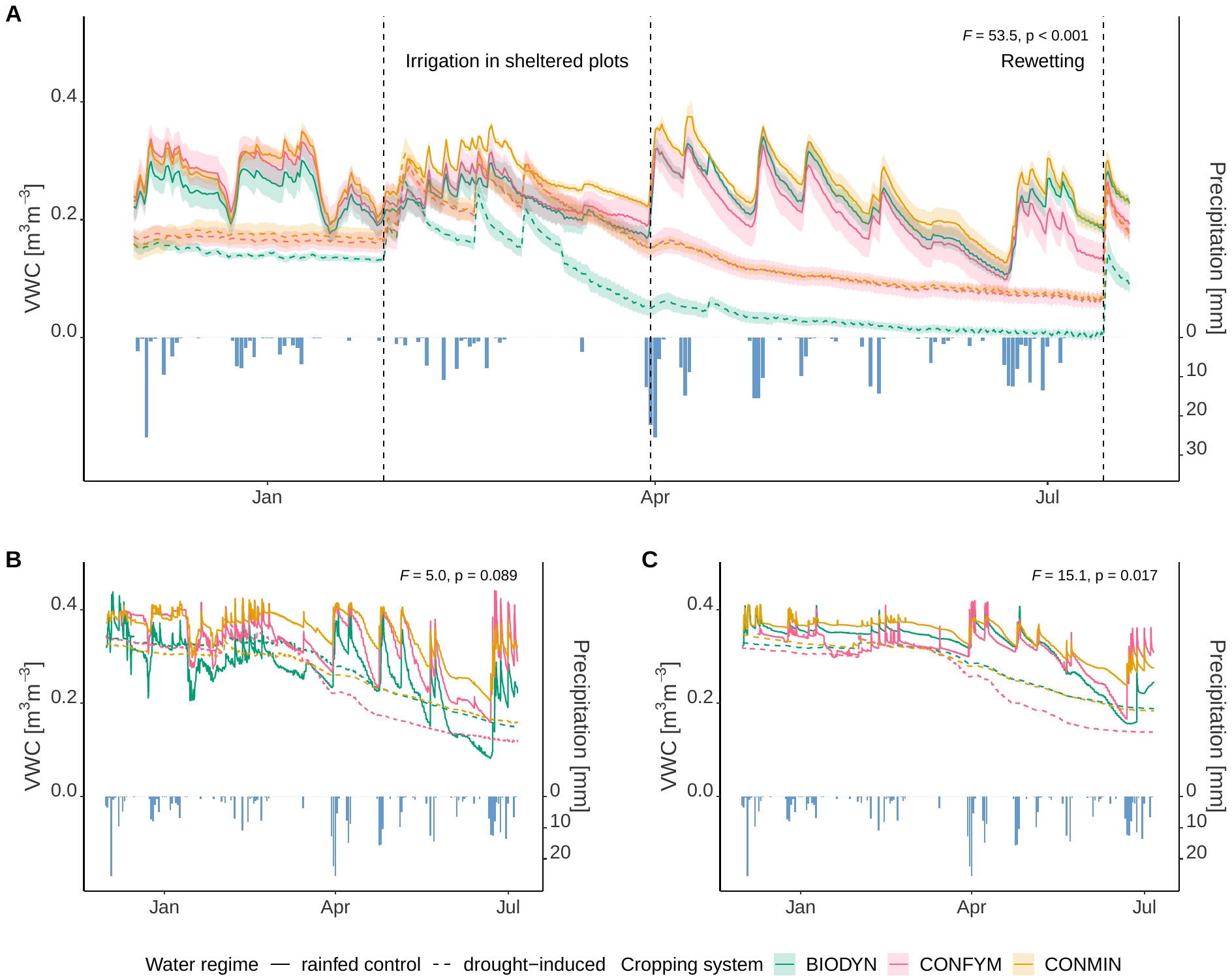

Supplementary Figure 2: Volumetric soil water content (VWC) in the sheltered and control plots of the three cropping systems. VWC was measured with TOMST sensors in all plots to a depth of 15cm and shown as means and errors deviations of four replications (n=4) (A), and with TDR sensors in three cropping systems x two water regimes x two depths at 5cm (B) and 20cm (C) depth. Bars in the plot represent the total daily precipitation in the rainfed control plots. TOMST and TDR measurements lasted until two weeks after rewetting and until wheat harvest, respectively. Significant differences between water regimes over the drought period (1 April to 14 July 2022) assessed by ANOVA adjusted for repeated measures are given as p and F-values.

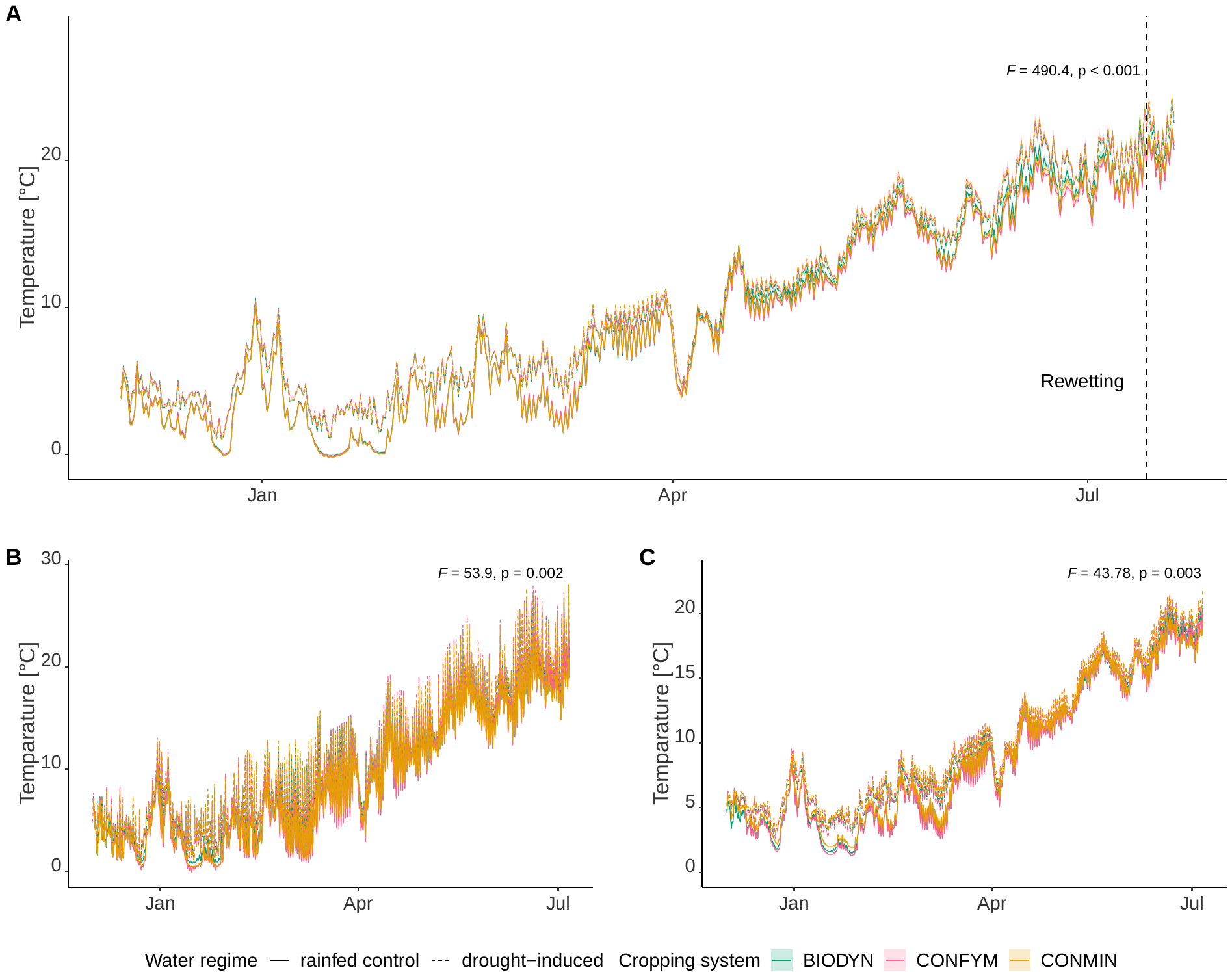

Supplementary Figure 3: Soil temperature in the sheltered and control plots of the three cropping systems. Soil temperature was measured with TOMST sensors in all replications to a depth of 6cm and shown as means and standard errors of four replications (A), and with TDR sensors in three cropping systems x two water regimes x two depths at 5cm (B) and 20cm (C) depth total. TOMST and TDR measurements lasted until two weeks after rewetting and until wheat harvest, respectively. Significant differences between water regimes assessed by ANOVA adjusted for repeated measures are given as p and F-values.

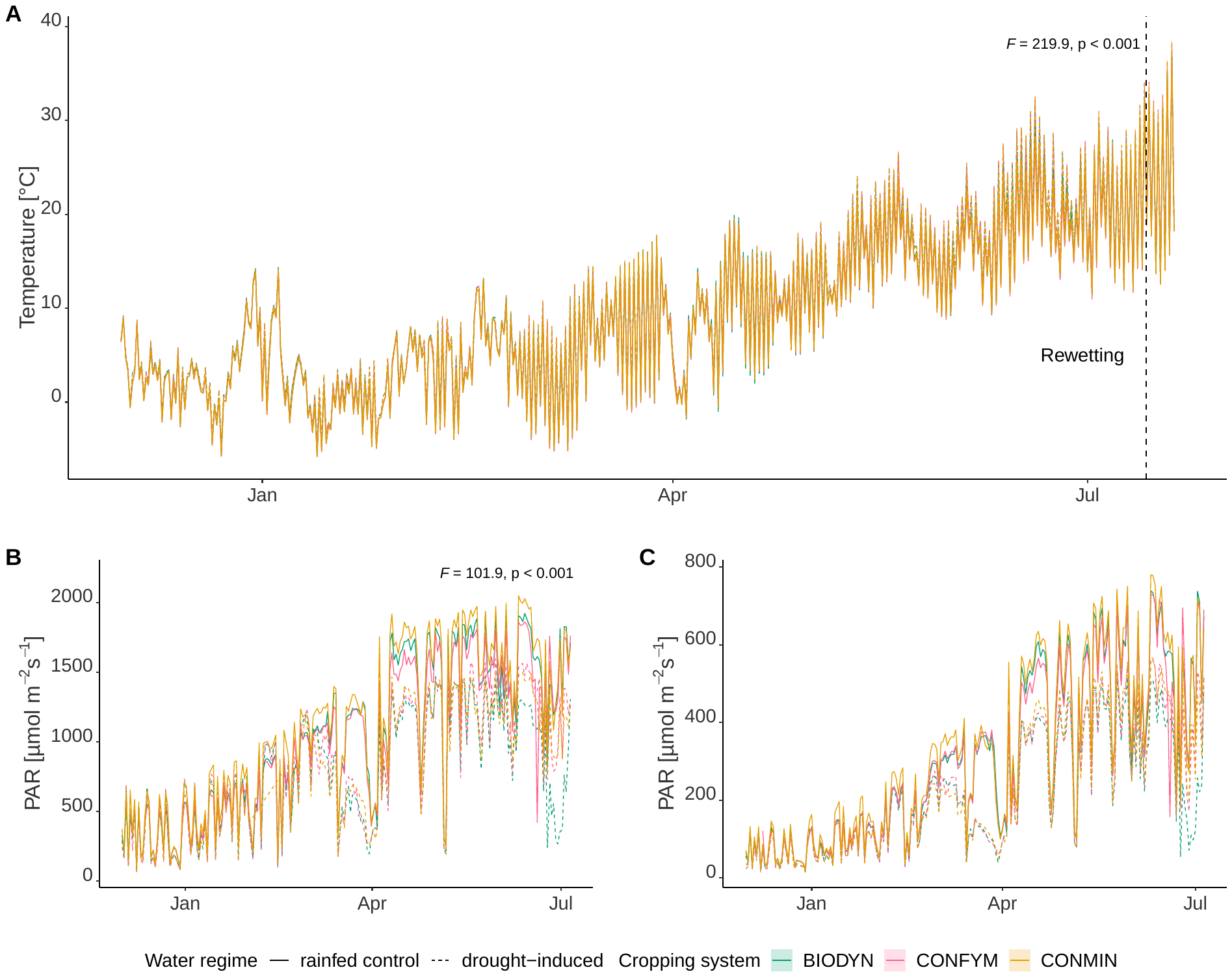

Supplementary Figure 4: Air temperature and photosynthetic active radiation (PAR) in the sheltered and control plots of the three cropping systems. Air temperature was measured with TOMST sensors at 12cm above the ground in all replications and is shown as means and standard errors of four replications (A), PAR was measured with PAR Photon Flux Sensors in three cropping systems x two water regimes (6 measurements totally) and provided as maximal daily PAR (B) and mean daily PAR (C). TOMST and PAR measurements lasted until two weeks after rewetting and until wheat harvest, respectively. Significant differences between water regimes assessed by ANOVA adjusted for repeated measures are given as p and F-values.

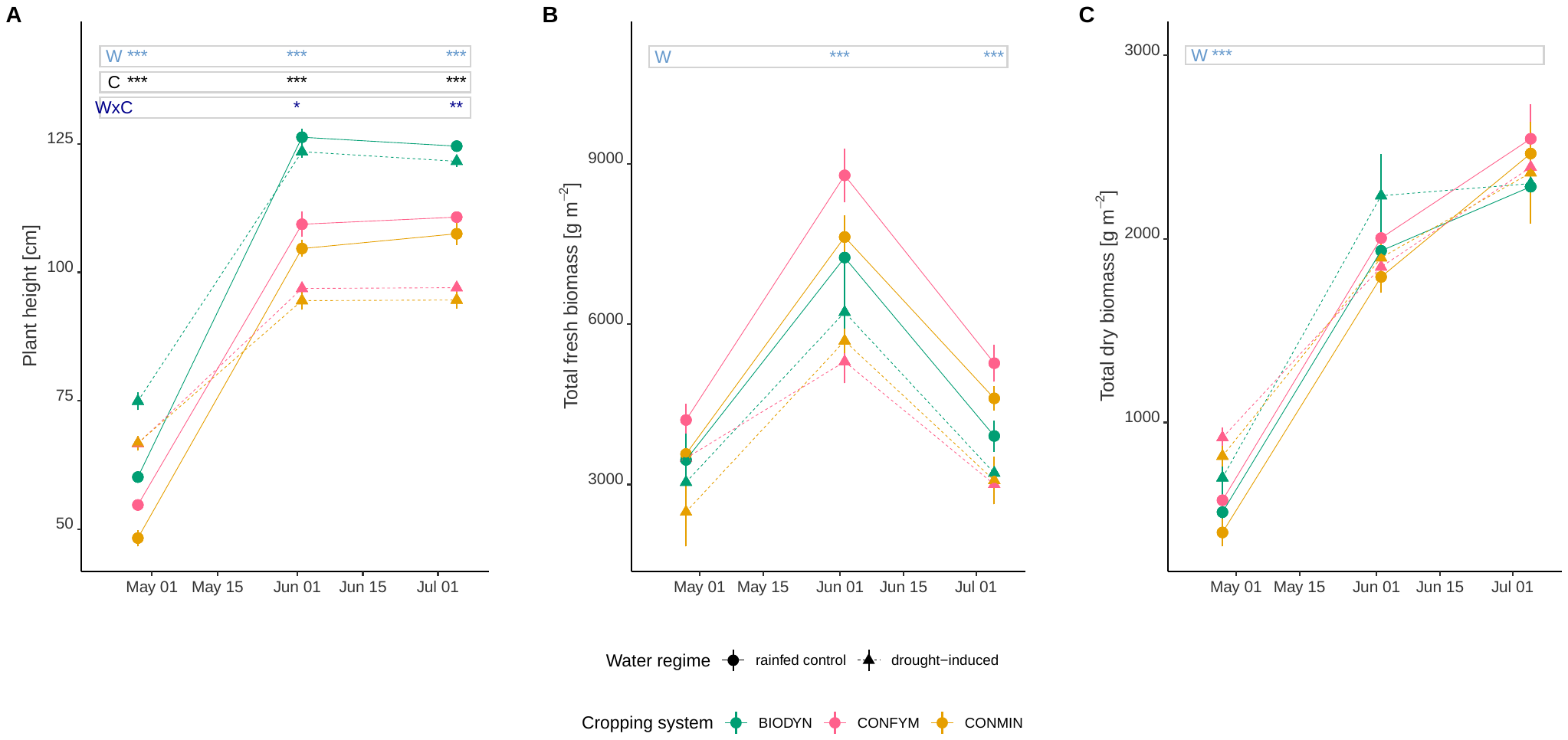

Supplementary Figure 5: Plant height (A), total fresh (B) and dry biomass (C) in the sheltered and control plots. Plant height. fresh and dry biomass was measured at three sampling dates (stem elongation, flowering, ripening). Mean values and standard errors (n=4) are provided. Significant differences between water regimes (W), cropping systems (C), and their interaction for each sampling date assessed by ANOVA are indicated with asterisks (* p < 0.05, ** p < 0.01, *** p < 0.001).

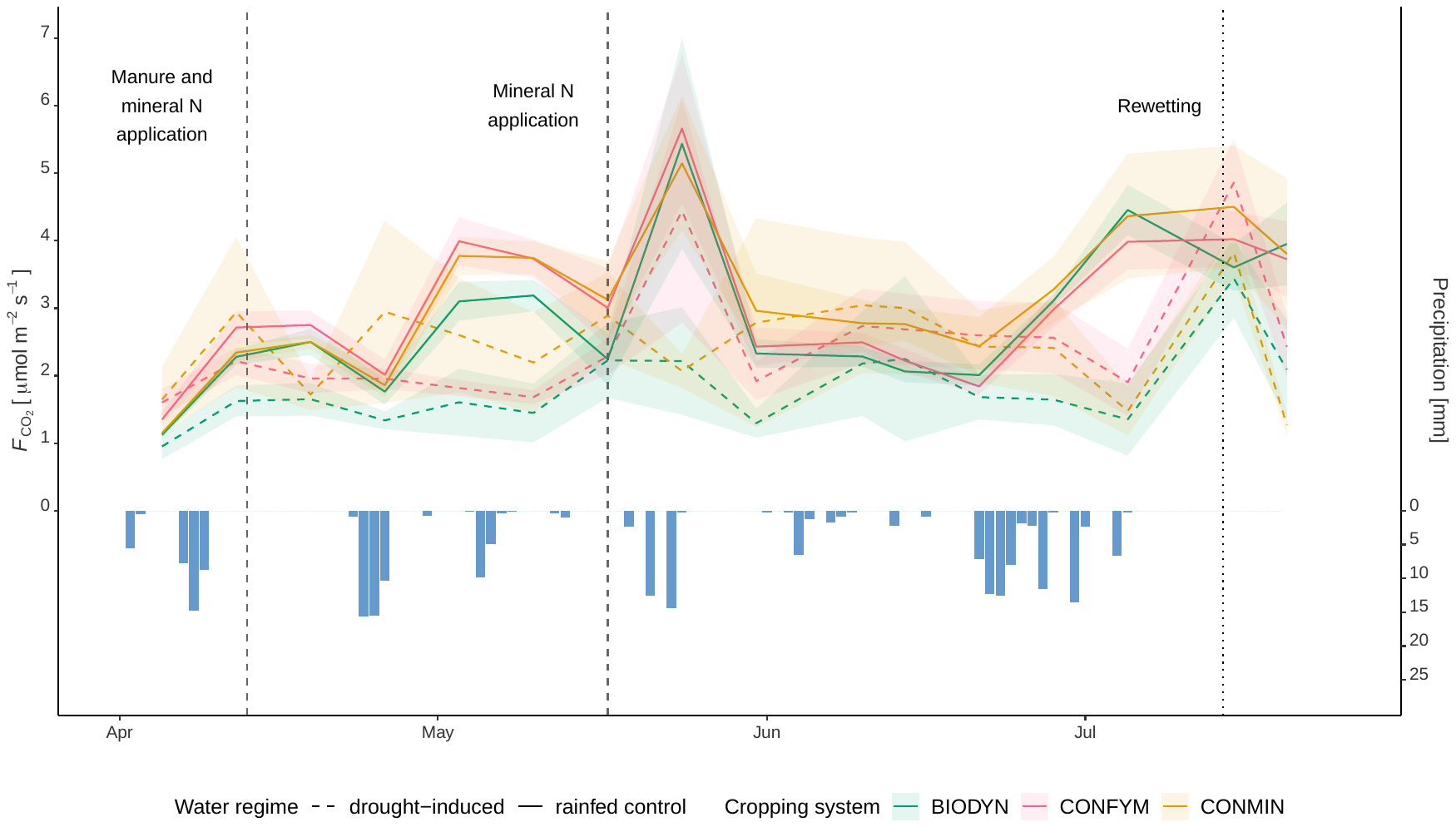

Supplementary Figure 6: Soil respiration in the sheltered and control plots. Soil respiration was weekly measured in-situ over the wheat vegetation period up to one week after rewetting. Mean values and standard erorrs are provided (n=4). Rewetting, manure, and mineral nitrogen (N) applications are indicated by vertical dashed lines. Blue bars represent the total daily precipitation for rainfed control plots.

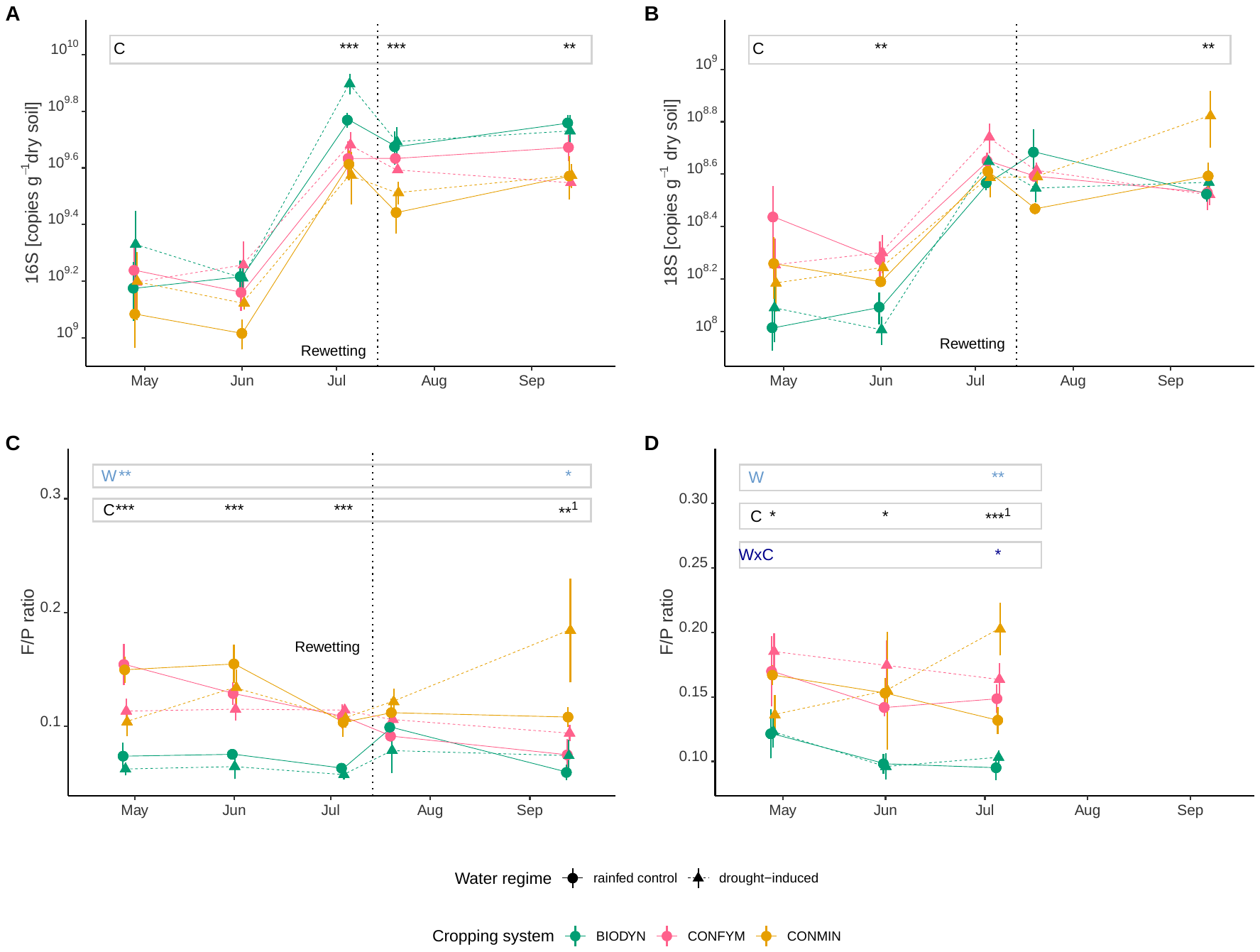

Supplementary Figure 7: Prokaryotic and fungal rRNA gene copy number and fungi: prokaryotes ratio in sheltered and control plots. Prokaryotic 16S (A) and fungal 18S (B) rRNA gene copy numbers measured in bulk soil as well as fungal to prokaryotic ratio (F/P ratio) in bulk soil (C) and rhizosphere (D) for each water regime and cropping system. Mean values and standard errors are provided (n=4). Significant differences between water regime (W), cropping systems (C), and their interaction for each sampling date as assessed by PERMANOVA based on Euclidean distances are indicated with asterisks (* p < 0.05, ** p < 0.01, *** p < 0.001). Heteroscedasticities are indicated as superscript ^1^.

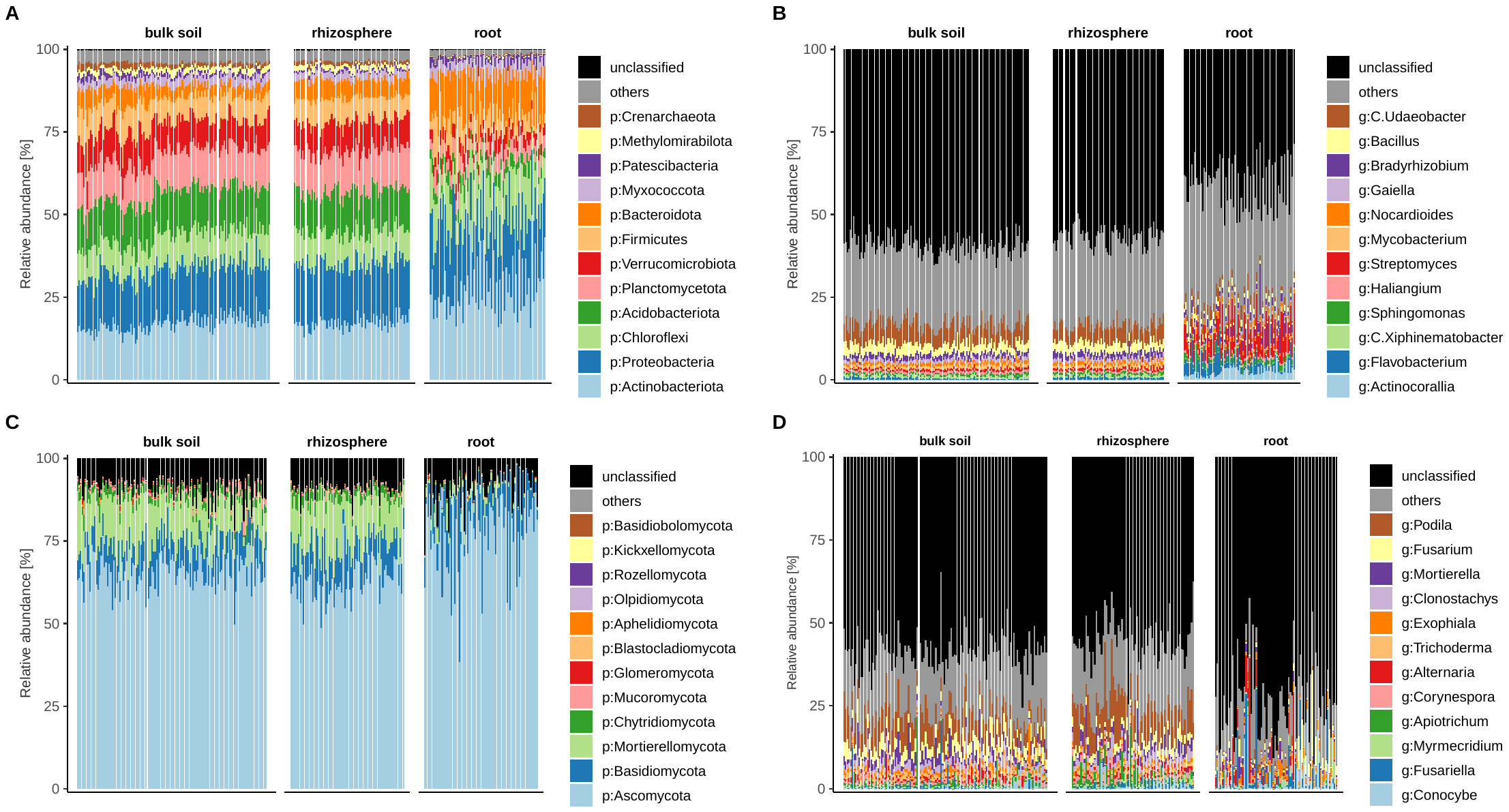

Supplementary Figure 8: Relative abundance of ASVs aggregated at phylum and genus level in the three investigated compartments, i.e. bulk soil, rhizosphere, and root. Bars show mean relative abundances per sample at phylum (A) and genus (B) level for prokaryotes, and at phylum (C) and genus (D) level for fungi.

Supplementary Figure 9: Principal coordinate analysis (PCO) ordinations of prokaryotic and fungal communities based on Bray-Curtis dissimilarities derived from ASV counts. PCOs are shown for prokaryotes in bulk soil (A), rhizosphere (C), and root (E), and for fungi in bulk soil (B), rhizosphere (D), and roots (F). The variance explained by each PCO axis are provided in parentheses.

Supplementary Figure 10: Effects of drought and sampling date on prokaryotic and fungal β-diversity as assessed by canonical analysis of principal coordinates (CAP) that maximizes discrimination between these factors. Panels represent differences in prokaryotic communities in bulk soil (A), rhizosphere (C), root (E), and fungal communities in bulk soil (B), rhizosphere (D), and root (F). The CAP overall reclassification rate in percentage, the Pillai’s trace statistics, and the statistical significance (p < 0.001 ***) are provided in each plot. Reclassification rates for each water regime at each sampling date are provided next to the respective ellipses. The amount of between-group variation of each CAP axis is provided in parentheses.

Supplementary Figure 11: Effects of drought and sampling date (drought, first and second resilience) on prokaryotic and fungal β-diversity during drought and after rewetting as assessed by canonical analysis of principal coordinates (CAP) that maximizes discrimination between these factors. The CAP overall reclassification rate in percentage, the Pillai’s trace statistics, and the statistical significance (p < 0.001 ***) are provided in each plot. Reclassification rates for each water regime at each sampling date are provided next to the respective ellipses. Panels represent differences in prokaryotic communities in bulk soil in CONMIN (A), CONFYM (C), BIODYN (E) and fungal communities in bulk soil in CONMIN (B), CONFYM (D), BIODYN (F). The amount of between-group variation of each CAP axis is provided in parentheses.

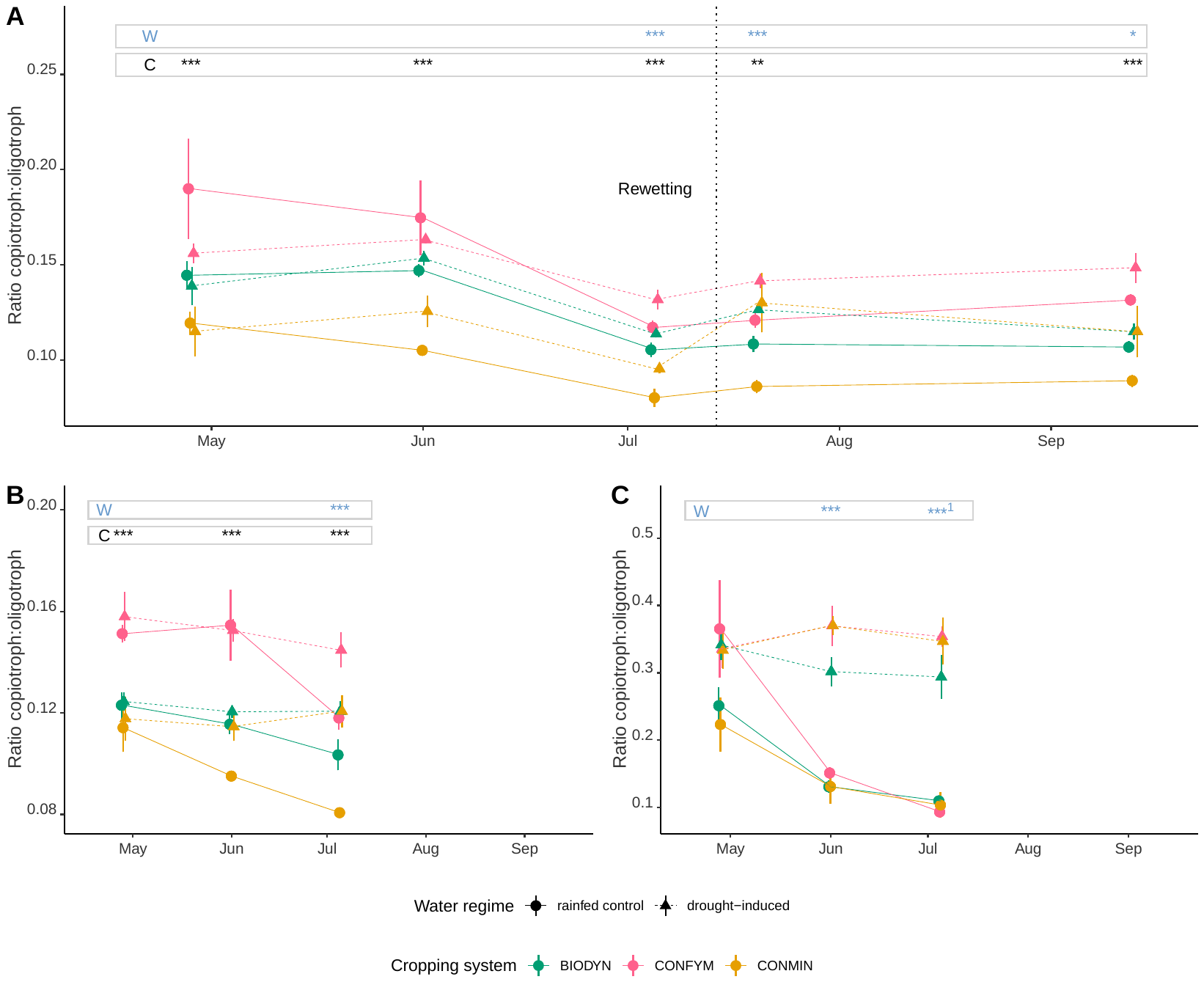

Supplementary Figure 12: Ratio of copiotrophs to oligotrophs in sheltered and control plots. Ratio was assessed based on estimated rRNA gene copy numbers of prokaryotic ASVs for the water regime and cropping systems using the thresholds of 5 for differentiating copiotrophic against oligotrophic taxa. Mean values and standard errors are provided (n=4) in A) bulk soil, B) rhizosphere, and C) root. Significant differences between water regime (W) and cropping systems (C) for each sampling date as assessed by PERMANOVA based on Euclidean distances are indicated with asterisks (* p < 0.05, ** p < 0.01, *** p < 0.001). Heteroscedasticities are indicated as superscript ^1^.
